# Immunohistochemical and ultrastructural characterization of the inner ear epithelial cells of splitnose rockfish (*Sebastes diploproa*)

**DOI:** 10.1101/2023.09.22.558895

**Authors:** Garfield T. Kwan, Leonardo R. Andrade, Kaelan J. Prime, Martin Tresguerres

## Abstract

The inner ear of teleost fish regulates the ionic and acid-base chemistry and secretes the protein matrix of the endolymph to facilitate otolith biomineralization, which are used to maintain vestibular and auditory functions. The otolith is biomineralized in a concentric ring pattern corresponding to seasonal growth, and this CaCO_3_ polycrystal has become a vital aging and life-history tool for fishery managers, ecologists, and conservation biologists. Moreover, biomineralization patterns are sensitive to environmental variability including climate change, thereby threatening the accuracy and relevance of otolith-reliant toolkits. However, the cellular biology of the inner ear is poorly characterized, which is a hurdle for a mechanistic understanding of the underlying processes. This study provides a systematic characterization of the cell types in the inner ear of splitnose rockfish (*Sebastes diploproa*). Scanning electron microscopy revealed the apical morphologies of the six inner ear cell types. Additionally, immunostaining and confocal microscopy characterized the expression and subcellular localization of the proteins Na^+^/K^+^-ATPase, carbonic anhydrase, V-type H^+^-ATPase, Na^+^-K^+^-2Cl^-^-Co-Transporter, Otolith Matrix Protein 1, and Otolin-1 in six inner ear cell types bordering the endolymph. This fundamental cytological characterization of the rockfish inner ear epithelium illustrates the intricate physiological processes involved in otolith biomineralization, and highlights how greater mechanistic understanding is necessary to predict their multi-stressor responses to future climate change.

## Introduction

The inner ear of teleost fishes regulates the ionic and acid-base chemistry of the endolymph and secretes the proteins necessary for otolith biomineralization (1). Each inner ear contains three CaCO_3_ otoliths called the sagittae, lapilli, and asteriscii, and each of these polycrystals serves as an inertial mass on adjacent sensory neurons that convey that information to the brain to maintain proper vestibular and auditory functions. Otoliths grow continuously throughout the fish’s life, so the inner ear must in turn expand to accommodate the larger polycrystal while maintaining proper homeostatic and sensory function. The inner ear’s capacity to regulate endolymph chemistry is consistently robust, and the resulting otolith biomineralization patterns have been utilized to derive a myriad of chronological information for over half a century (1, 3).

Otoliths are biomineralized in a concentric ring pattern that follows seasonal growth, thereby forming annual rings called “annuli” that serve as a proxy for fish age (4, 5). Fishery managers incorporate this information into age-structured models to estimate growth rate, maturity age, recruitment, mortality, and overall population biomass (6, 7). These otolith-reliant age-structured models are used to make informed decisions on catch limits to sustainably regulate fish stocks around the world, thereby preserving both aquatic ecosystems and economies. In addition, otolith microchemistry has become an important tool to derive life history information. The analysis of trace elements and isotopes trapped within the otolith have been used to retroactively determine a variety of metabolic and environmental factors experienced by the fish such as metabolic rate (8, 9), temperature (10), diet (11), hypoxia (12), and salinity (13). These otolith-derived life-history information is particularly useful for agencies that manage endangered and migratory species as they explore methods to determine recruitment and dispersal (14), stock origin, migration pattern, and pollution exposure (15–17).

Past studies on the inner ear has described the organ in four regions: the macula, meshwork area, intermediate area, and patches area (18, 19), with the otolith resting on a gelatinous otolithic membrane that covers much of the macula (20–22) (Figure 1). The otolithic membrane is a thin gelatinous membrane separating the macula from the otolith (Figure 1), and each sockets usually contains the hair bundles from a single HC (20, 21). The otolithic membrane protects the HCs from being crushed by the overlying otolith as well as in distributing the inertial movement across the hair cells (20). The center of the macula is comprised of HCs and SCs, and the periphery of the macula contains GCs (Figure 1). The HCs are known to have basolateral NKA and cytoplasmic VHA in its basolateral soma (23, 24), tubulin-rich hair bundle (25), and rely on K^+^ inflow as the driving force for its electrochemical potential similar to mammalian HCs (26, 27). The surrounding SCs are thought to aid in secreting the matrix proteins, and anchoring the filamentous material with cell boundary microvilli as proposed for inner ear systems of aves and amphibians (20, 28, 29). The GC has an extensive endoplasmic reticulum and numerous secretion granules (18) to synthesize and secrete proteins such as Otolin-1 to chelate with Ca^2+^ ions to promote biomineralization (30–35). The meshwork area is known to have T1-Is and T2-Is (Figure 1). The T1-Is are known to be mitochondrion-, NKA- and NKCC-rich (24, 36–38), have short thick microvilli (18), and thought to be responsible maintaining the ionic and acid-base parameters of the endolymph (19, 24, 37, 39, 40). T1-Is are interconnected and together form intracytoplasmic membrane systems, and their abundance decreases with distance away from the macula (18, 38). In contrast, the T2-Is are known to contain CA (41, 42) and VHA (24, 37), and they become the more common with distance away from the macula. In fact, the intermediate area is dominated by T2-Is (Figure 1). Finally, the patches area predominantly contains SQs (Figure 1). The SQs have been called “small ionocytes”, and they are generally very “flattened” with highly infolded lateral plasma membranes and elongated mitochondria (18, 38). SQs also strongly express CA (41, 42) and OMP-1 (43), and their proposed role(s) include the secretion of ions and bicarbonate (41, 42) along with OMP-1 to facilitate biomineralization on the distal side of the otolith (19, 40, 44). Notably, the overwhelming majority of our current inner ear cell characterizations are derived from freshwater species such as the zebrafish (*Danio rerio*) and rainbow trout (*Oncorhynchus mykiss*); this greatly contrasts with the application of otolith-based tools, which is heavily used to regulate marine fisheries.

**Figure 1:**
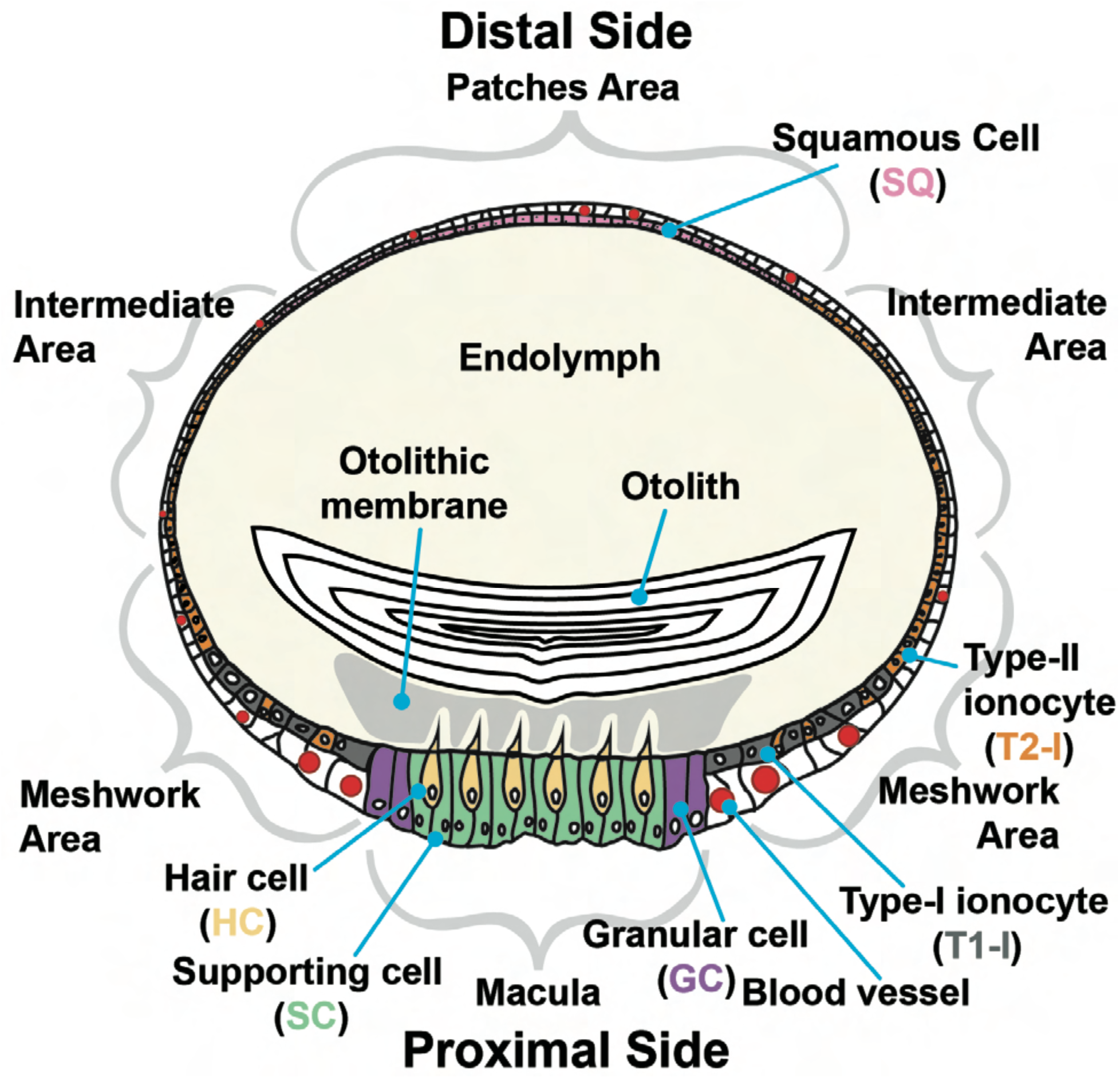
Schematic of the inner ear organ. The inner ear is categorized into four general areas: the macula, meshwork area, intermediate area, and patches area. The macula contains the hair cell (HC), supporting cell (SC), and granular cell (GC). The meshwork area contains the Type-I ionocyte (T1-I) and Type-II ionocyte (T2-I). The intermediate area only contains T2-I, and the patches area contains only squamous cell (SQ). The macula and meshwork area are considered to be on the proximal side of the inner ear, and the intermediate and distal areas are considered to be on the distal side of the inner ear. The organ is interwoven with blood vessels, most of which are on the proximal side of the inner ear.

Recent research has suggested that near-future climate change scenarios could significantly alter otolith biomineralization patterns thereby threatening the many otolith-reliant analytical techniques that researchers have relied upon for the past half century. In general, exposure to elevated CO_2_ (45, 46) and warming (47, 48) accelerates otolith growth, whereas hypoxia hinders biomineralization (49–51) and can even generate false annual rings known as “checks” (49, 50). Indeed, inaccurate otolith-based aging and data misinterpretation *can* and *have* led to unsustainable fisheries, for example, those of the orange roughy (*Hoplostethus atlanticus*) in New Zealand (52) and the walleye pollock (*Gadus chalcogrammus*) in Bering Sea (53). Despite the need to understand their cell-specific differences in morphology, protein localization and abundance, the characterization of inner ear cells and their contribution to otolith biomineralization have received scant attention (1), and is recognized as a critical knowledge gap by the international sclerochronology community (2).

Since otoliths are formed inside the inner ear endolymph and are not in direct contact with the water, the altered biomineralization pattern and the accumulation of trace element and isotopes within the polycrystal necessitates a degree of biological regulation by the inner ear epithelial cells. Indeed, the endolymph has higher average pH, [HCO_3_−], [CO_3_^2−^], total CO_2_, and [K^+^] than the blood plasma (24, 39, 40, 54, 55). Moreover, the endolymph in the proximal side of the inner ear organ (facing the macula) has much lower [K^+^] and pH than the distal side (facing the patches area) (19, 40), a chemical heterogeneity that is most likely generated by the activity of the inner ear epithelial cells in each region. Further complicating matters is that the inner ear is known to contain at least six types of cells (18, 19) (Figure 1), each with their own specific morphology and protein localization/abundance to fulfill their function(s).

Previous studies have examined the ultrastructure of the inner ear epithelial cells using transmission electron microscopy (TEM) and the expression of individual proteins in specific cell types using immunohistochemistry and fluorescence microscopy on tissue sections, scanning electron microscopy (SEM), (18, 20, 21, 36–38, 41, 43, 56). Here, we combined multiplex immunohistochemistry and SEM on whole-mounted inner ear tissue to provide a comprehensive immunological and ultrastructural characterization of the six cell types lining the endolymph. This approach allowed us to generate unprecedent three-dimensional reconstructions of the expression patterns of NKA, CA, VHA, NKCC, OMP-1 and Otolin-1 in the six inner ear epithelial cell types that border the endolymph, together with matching visualizations of their apical morphology. Altogether, our results highlight the high cytological complexity of the fish inner ear epithelium and provides critical foundational knowledge to help decode the mechanisms responsible for otolith biomineralization and their regulation.

## Methods

### Tissue Sampling

Young-of-year splitnose rockfish (*Sebastes diploproa*) were caught from drifting kelp paddies off the shores of La Jolla and raised in the Hubbs Experimental Aquarium at the Scripps Institution of Oceanography, University of California San Diego (La Jolla, USA). The animals were raised in a flow-through system with seawater continuously pumped from the Scripps Coastal Reserve for >2 years, and were fed EWOS food pellet (Cargill Incorporated, Minneapolis, MN, USA). In accordance with protocol S10320 of the University of California, San Diego Institutional of Animal Care and Use Committee, fish were euthanized with an overdose of anesthetic (tricaine mesylate 0.2 g/L buffered with NaOH) followed by spinal pithing before inner ear tissue sampling. In total, 24 splitnose rockfish (total length: 11.88 ± 0.29 cm; weight: 42.86 ± 2.79) were analyzed.

### Antibodies

NKA was immunodetected using the mouse monoclonal anti-NKA antibody α5 (57) purchased from the Developmental Studies Hybridoma Bank (DSHB, The University of Iowa, Iowa City, IA, USA). VHA was immunodetected using custom-made rabbit polyclonal antibodies against a highly conserved epitope within subunit B (epitope: AREEVPGRRGFPGY; GenScript, Piscataway, USA). Both NKA and VHA antibodies have been validated in the inner ear of splitnose rockfish (24) and Pacific chub mackerel (*Scomber japonicus*) (37). CA was immunodetected using rabbit polyclonal antibodies against human CA II (Rockland Inc., Gilbertsville, USA), and this antibody has been validated in the inner ear of Masu Salmon (*Oncorhynchus masou*) (42) and Pacific chub mackerel (37). NKCC was immunodetected using the anti-NKCC antibody T4 (58) obtained from DSHB, and this antibody has been validated in the inner ear of zebrafish (59) and Pacific chub mackerel (37). OMP-1 and Otolin-1 were immunodetected using antibodies generously provided by Dr. Emi Murayama (Institut Pasteur, Paris, France), which were previously validated in rainbow trout inner ear (43). The secondary antibodies were goat anti-mouse IgG-HRP and goat anti-rabbit IgG-HRP conjugate (Bio-Rad, Hercules, CA, USA) for western blotting, and goat anti-mouse Alexa Fluor 546, goat anti-rabbit Alexa Fluor 488, and/or goat anti-rabbit Alexa Fluor 555 (Invitrogen, Grand Island, USA) for immunohistochemistry.

The specificity of NKA, VHA, CA, NKCC, OMP-1, and Otolin-1 antibodies within splitnose rockfish inner ear tissues was verified using western blotting (methods below). NKA, VHA, CA, and NKCC yielded bands matching their respective predicted size at ∼100, ∼55, ∼40, ∼210, ∼80 and ∼100 kDa (Supplementary Figure 1).

### Western Blotting

Frozen inner ear samples were immersed in liquid nitrogen, pulverized using a porcelain mortar and pestle, and submerged within an ice-cold, protease inhibiting buffer (250 mmol l^−1^ sucrose, 1 mmol l^−1^ EDTA, 30 mmol l^−1^ Tris, 10 mmol/L benzamidine hydrochloride hydrate, 1 mmol/L phenylmethanesulfonyl fluoride, 1 mmol/L dithiothreitol, pH 7.5). Samples were centrifuged at 3,000xg (10 min, 4°C) to remove debris, and the resulting supernatant was saved as the sample. Total protein concentration in the crude homogenate was determined by the Bradford assay (60). On the day of SDS-electrophoresis, the samples were mixed with an equal volume of 2x Laemmli buffer containing 10% β-mercaptoethanol, heated (70°C for 5 min) and loaded (10 µg per lane) onto a 7.5% polyacrylamide mini gel (Bio-Rad, Hercules, CA, USA). Proteins in the samples were separated by polyacrylamide gel electrophoresis at 200 V for 40 min, then transferred onto a polyvinylidene difluoride (PVDF) membrane using a wet transfer cell (Bio-Rad) at 100 mAmps (4°C overnight). On the following day, PVDF membranes were incubated in tris-buffered saline with 1% tween (TBS-T) with milk powder (0.1 g/mL) for 1 h, then incubated with primary antibody (a5: 10.5 ng/ml; H300: 100 ng/ml; VHA *b*-subunit: 1.5 µg/ml; CA II antibody: 8 µg/ml; T4: 10.4 ng/ml; OMP-1: 1:1000 dilution; Otolin-1: 1:1000 dilution) at 4°C overnight. On the following day, PVDF membranes were washed in TBS-T (3 x 10 min), incubated in blocking buffer with the appropriate secondary antibodies (1:10,000 in blocking buffer) for 1 h, and washed again in TBS-T (3 x 10 min). Bands were made visible through addition of ECL Prime Western Blotting Detection Reagent (GE Healthcare, Waukesha, WI) and imaged and quantified in the Universal III Hood (BioRad) using Image Lab software (version 6.0.1; BioRad).

### Immunostaining

Inner ears were fixed in 4% paraformaldehyde in phosphate buffer saline (PBS) at 4°C for 8 hours, incubated in 50% ethanol for 8 hours, and stored in 70% ethanol. On the day of analysis, samples were re-hydrated in PBS for 10 minutes. Native autofluorescence was reduced using sodium borohydride (1 mg/mL) in ice cold PBS (6 times; 10 minutes each) as demonstrated in (61). Samples were then washed in PBS + 0.1% tween (PBS-T) at RT for 5 minutes, incubated in blocking buffer (PBS-T, 0.02% normal goat serum, 0.0002% keyhole limpet hemocyanin) at RT for one hour, and with the primary antibodies (a5: 42 ng/ml; VHA subunit B: 6 µg/ml; CA II antibody: 160 µg/ml; T4: 104 ng/ml; OMP-1: 1:500; Otolin-1: 1:500) in blocking buffer and kept in a humid chamber at RT overnight. On the following day, samples were washed in PBS-T (3 times; 10 minutes each) and incubated with the appropriate anti-rabbit or anti-mouse fluorescent secondary antibodies (1:1,000) and nuclear stain DAPI (1 µg/mL) at RT for 1 hour. Samples were washed in PBS-T (three times; 10 minutes each), then whole-mounted and flattened onto depressed glass slide fitted with a glass cover slip (No. 1.5, 0.17 mm). Samples were Z-stacked imaged (Zeiss ZEN 2.6 blue edition software) using an inverted confocal microscope (Zeiss AxioObserver Z1 with LSM800; Oberkochen, Germany) at low (20x objective: Plan-Apochromat 20x/0.8 M27), high magnifications (40x objective: Zeiss LD LCI Plan-Apochromat 40x/1.2 lmmKorr DICM27 objective), and with Zeiss Airyscan super-resolution detector. Comparison of relative fluorescent signal across the different cell types were assessed in images visualized as maximum intensity projection or orthogonal cuts across the X-Z and Y-Z planes. Each of the six antibodies was immunotargeted in at least three rockfish.

### Scanning Electron Microscopy

Inner ears were dissected and immediately immersed in 4% paraformaldehyde and 2.5% glutaraldehyde in 0.2 M sodium cacodylate buffer at RT for 2 hours. Samples were washed in the same buffer at RT (three times; 15 minutes each), post-fixed in 1% osmium tetroxide (Electron Microscopy Science, Hatfield, PA, USA) at RT for 1 hour in dark, washed with ultrapure water (three times; 15 minutes each), incubated with 1% tannic acid (Electron Microscopy Science) at RT for 1 hour, washed again with ultrapure water (three times; 15 minutes each), and incubated a second time in 1% osmium tetroxide at RT for 1 hour. After yet another round of ultrapure water washes (three times; 15 minutes each), samples were dehydrated in an ethanol series until absolute (3x), critical-point dried (Bal-Tec CPD-030), sputter-coated with 10 nm of gold-palladium (Leica EM SCD 500) and observed in Zeiss Sigma VP operated at 5 kV. Each cell type was examined in at least three rockfish.

### Microvilli Measurements and Statistical Analysis

Microvilli length and width (N=20 microvilli per cell type) on the apical surface of T1-Is and T2-Is were measured with FIJI (version 2.1.0), then analyzed with one-way analysis of variance (ANOVA) using R (version 4.0.3) (62). The assumption of normality was checked using Shapiro-Wilks test, and an α of 0.05 was applied.

## Results

### Macula: Hair cells (HCs) and supporting cells (SCs)

SEM imaging shows that the hair bundle of each HC protrudes through a single opening in the gelatinous otolithic membrane (Figure 2A). Each HC hair bundle is composed of stereocilia arranged in a staircase pattern and one longer kinocilium at the top (Figures 2B, C). A circumstantial tear in the tissue revealed the soma of the HC, which has an ovoid shape (Figure 2C). An unobstructed, higher magnification view shows the presence of numerous pores and budding vesicles (Figure 2B, C), especially in the SCs (Figures 2B). Finally, microvilli are found on the SCs, but not on the HCs (Figure 2B).

**Figure 2:**
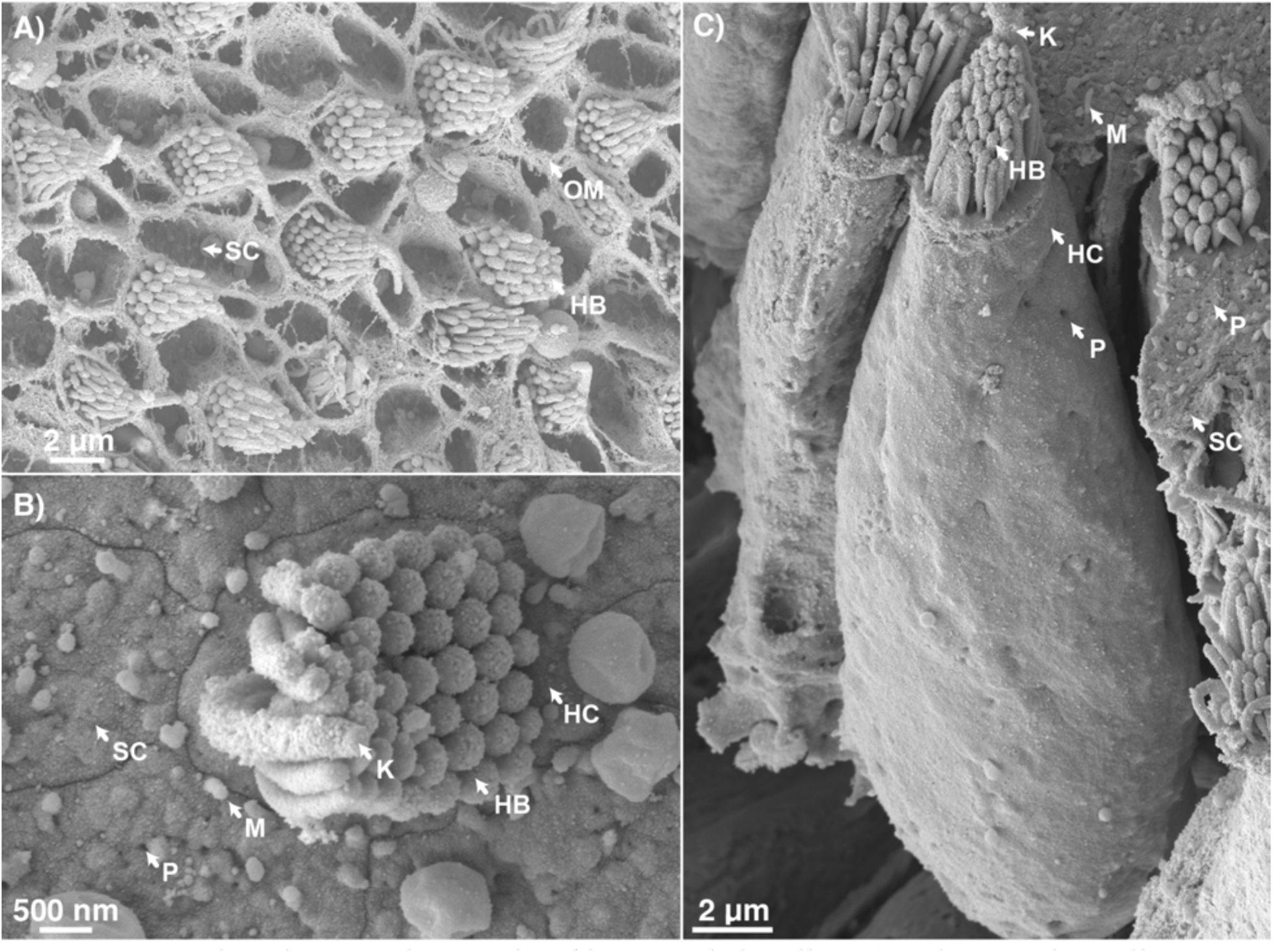
Scanning electron micrographs of inner ear hair cell (HC) and supporting cell (SC). A) The hair bundles (HB) of the HC protrudes through the otolith matrix (OM). B) When unobstructed, the pores (P) and microvilli (M) on the SC are visible. C) When viewed from the side, pores are also apparent on the HC’s basolateral membrane. Abbreviation: K=Kinocilium.

Three-dimensional reconstruction images of whole-mounted and immunostained inner ear tissue provide further insights about the inner ear epithelium structure and protein expression. Every HC has the nucleus basally located and its basolateral membrane expresses abundant NKA that outlines the ovoid shape of the soma (Figure 3; Supplemental Figure 2A). The HCs express CA at relatively low levels in their hair bundle (Figure 3A-D; Supplemental Figure 2A), and they abundantly express VHA throughout the cytoplasm (Figure 3E-H) and in their kinocilium of their tubulin-rich hair bundle (Supplemental Figure 2B). The HCs also has low expression of NKCC on its basolateral membrane (Figure 4), and occasionally express OMP-1 and Otolin-1 within their subapical area (Figure 5).

**Figure 3:**
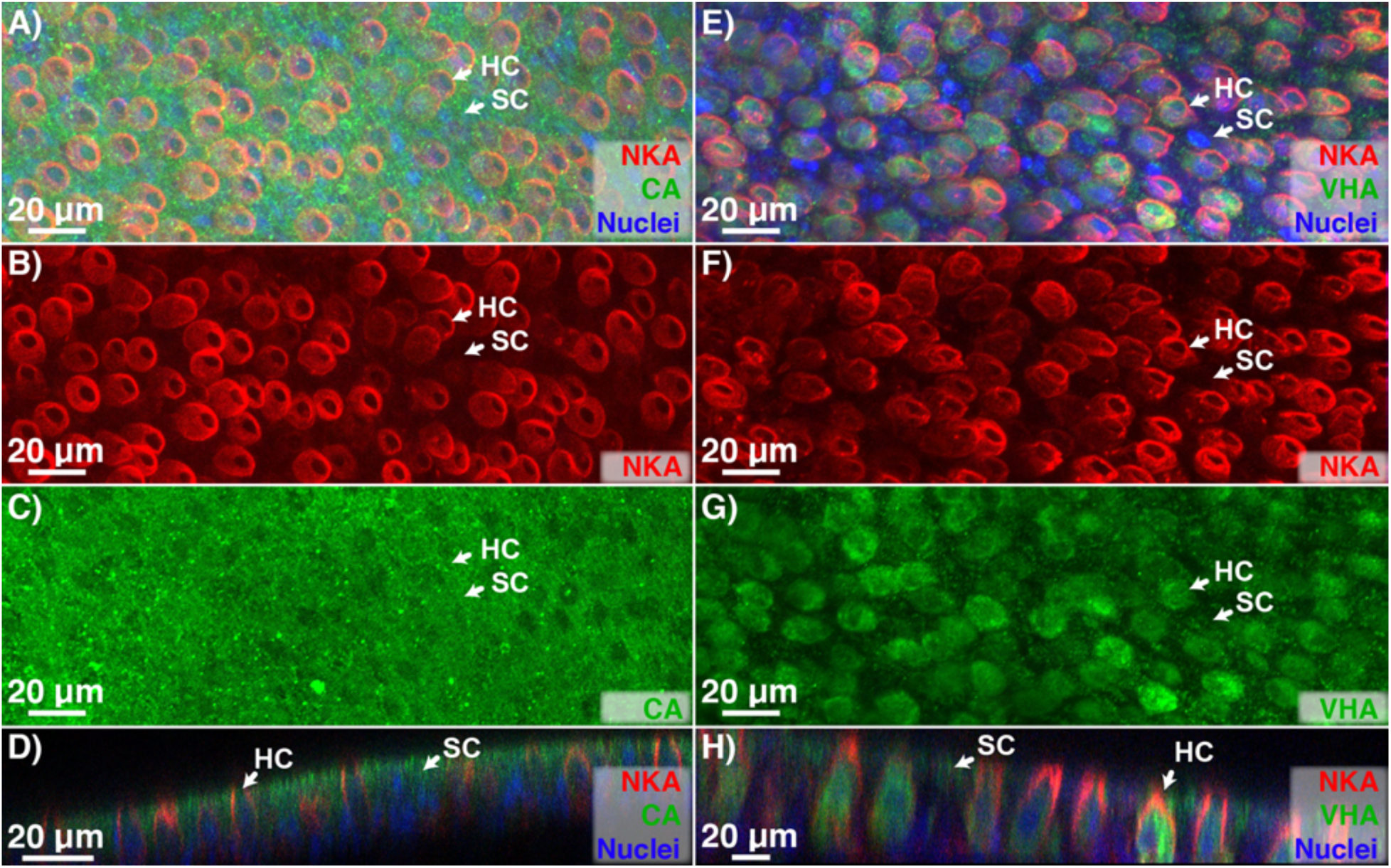
Immunolocalization of NKA, VHA, and CA in hair cell (HC) and supporting cell (SC). A, B) HC expresses NKA in the basolateral membrane, C, D) CA in their hair bundles (see supplemental figure 2), and E-H) VHA in their cytoplasm. A-D) The adjacent SC does not express NKA, but does express apical and subapical CA and VHA.

**Figure 4:**
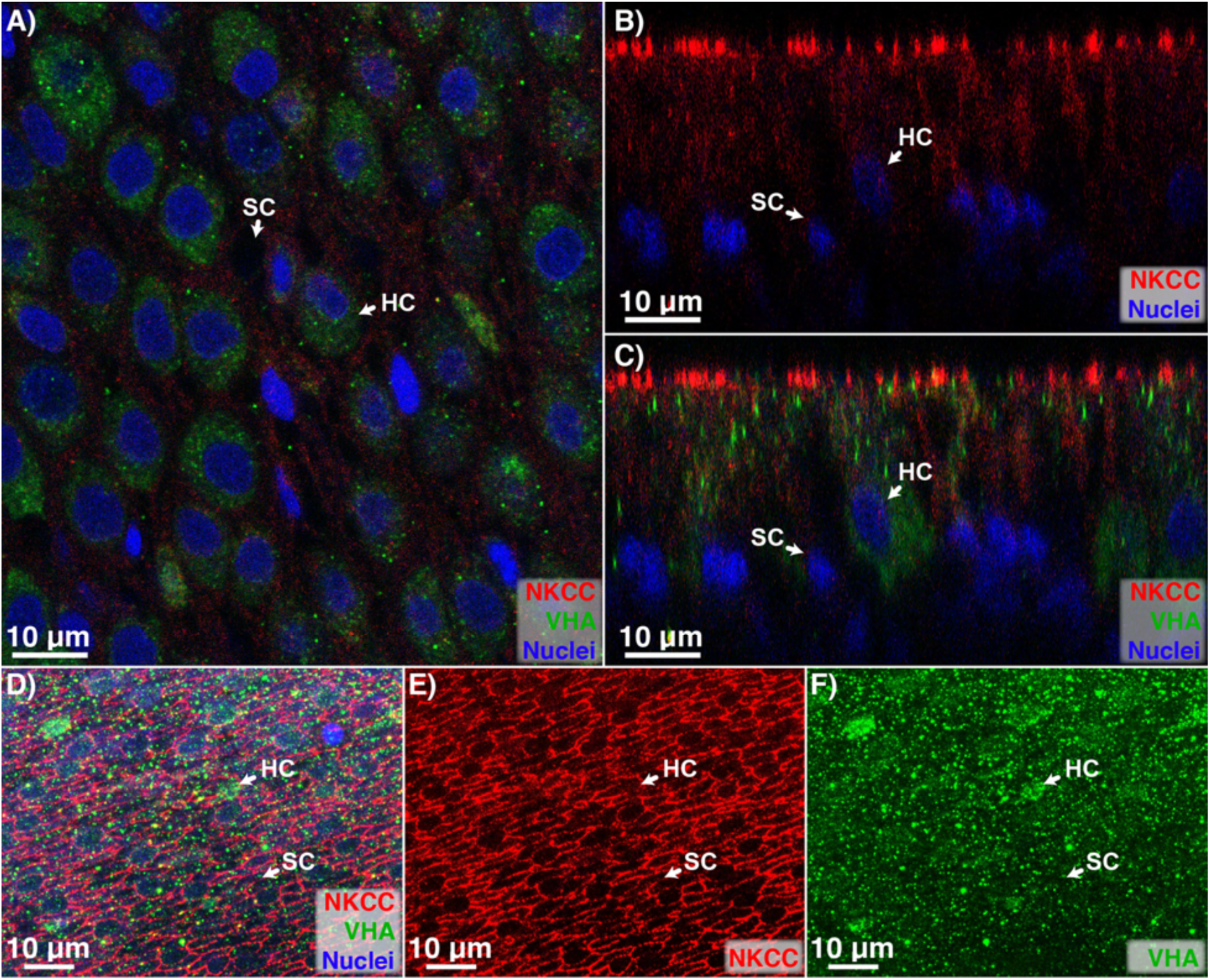
Immunolocalization of NKCC and VHA in hair cell (HC) and supporting cell (SC). A) The VHA-rich HC is surrounded by SC, which expresses B, C) basolateral NKCC as well as D-F) apical NKCC in a honeycomb pattern that resembles the overlying protein matrix.

**Figure 5:**
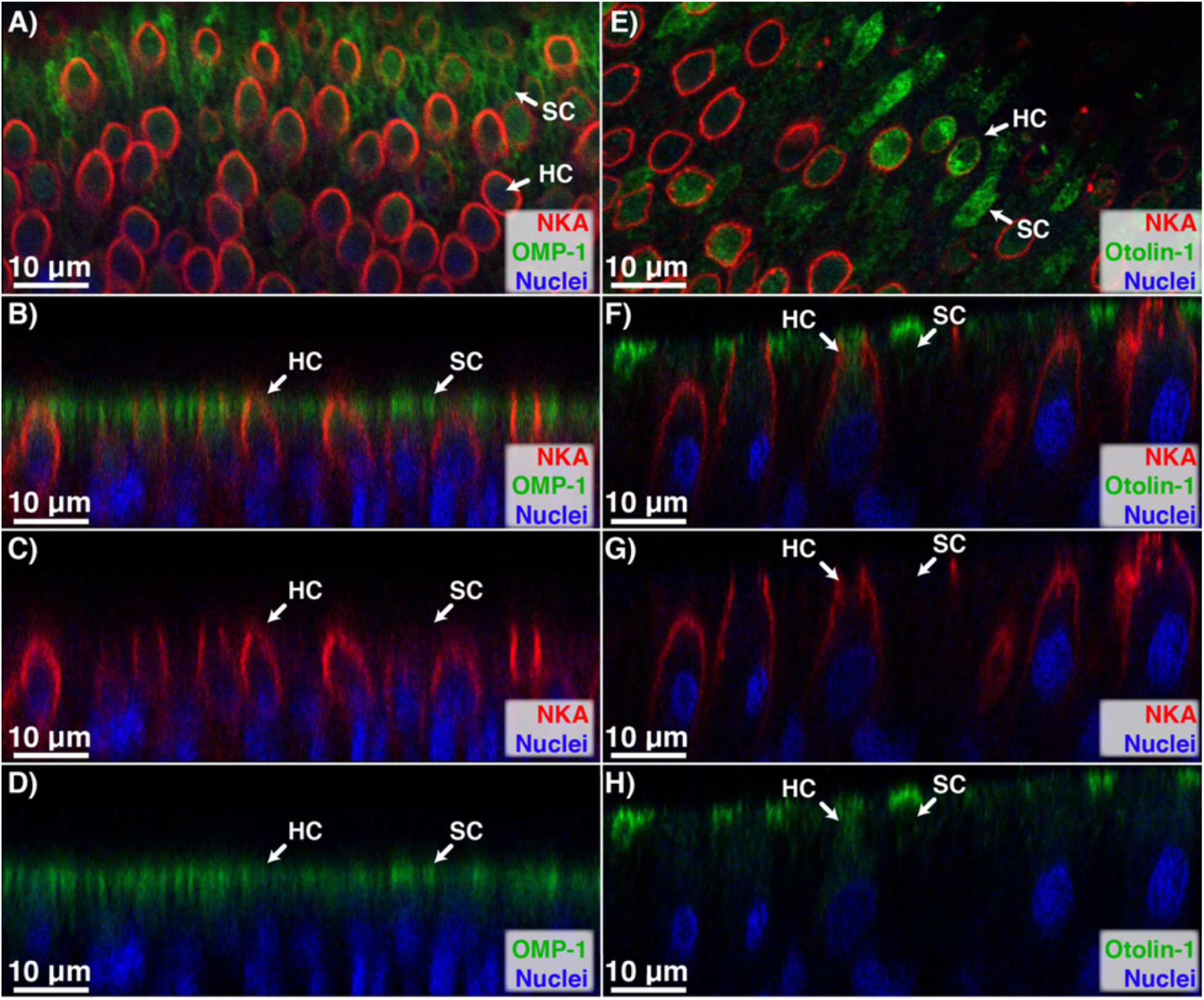
Immunolocalization of NKA, OMP-1, and Otolin-1 in hair cell (HC) and supporting cell (SC). The NKA-rich HC expresses some A-D) OMP-1 and E-H) Otolin-1 in its sub-apical area. In contrast, the SC expresses A-D) OMP-1 in a honeycomb pattern that resembles the overlying protein matrix, and E-F) otolin-1 in its sub-apical area.

The SCs extends past and envelope the basolateral membrane of the HCs, and the nucleus of the SC is located at the basal portion of the cell below that of the HCs (Figure 4B, C). Thus, a lateral view of three-dimensional reconstructions shows two parallel rows of nuclei that clearly distinguish HCs from SCs (Supplemental Figure 2C). The SCs do not show appreciable NKA signal (at least compared to the HCs); however, they express abundant CA throughout the cytoplasm and especially in the subapical area (Figure 3). Notably, a view from the top shows that the CA signal follows a honeycomb pattern resembling that of the SCs perimeter (Figure 3C). The SCs also express VHA in the subapical area; however, the signal is much dimmer than in the HCs (Figure 3G, H, Supplemental Figure 2C). The NKCC signal is detected on both the basolateral (Figure 4A-C, Supplemental Figure 2C) and apical side (Figure 4D, E; Supplemental Figure 2C); interestingly, the latter forms a honeycomb pattern resembling the extracellular otolithic membrane. The SCs express highly abundant OMP-1 and Otolin-1 in their subapical area (Figure 5), although the Otolin-1 signal is somewhat patchy (Figure 5H). Like CA and NKCC, OMP-1 signal also forms a honeycomb pattern (Figure 5A) whereas Otolin-1 appears throughout the apical surface of both HCs and SCs (Figure 5E).

### Macula: Granular cells (GCs)

The GCs line the periphery of the macula, separating the HCs and SCs from the T1-I and T2-I in the meshwork area. SEM imaging reveals the apical surface of the GCs has rhombic or pentagonal shape and contains numerous microvilli, pore-like structures, and budding vesicles (Figure 6A). In addition, small hair bundles are occasionally seen protruding in between the GC (Figure 6B-D), which reminisce incipient HCs.

**Figure 6:**
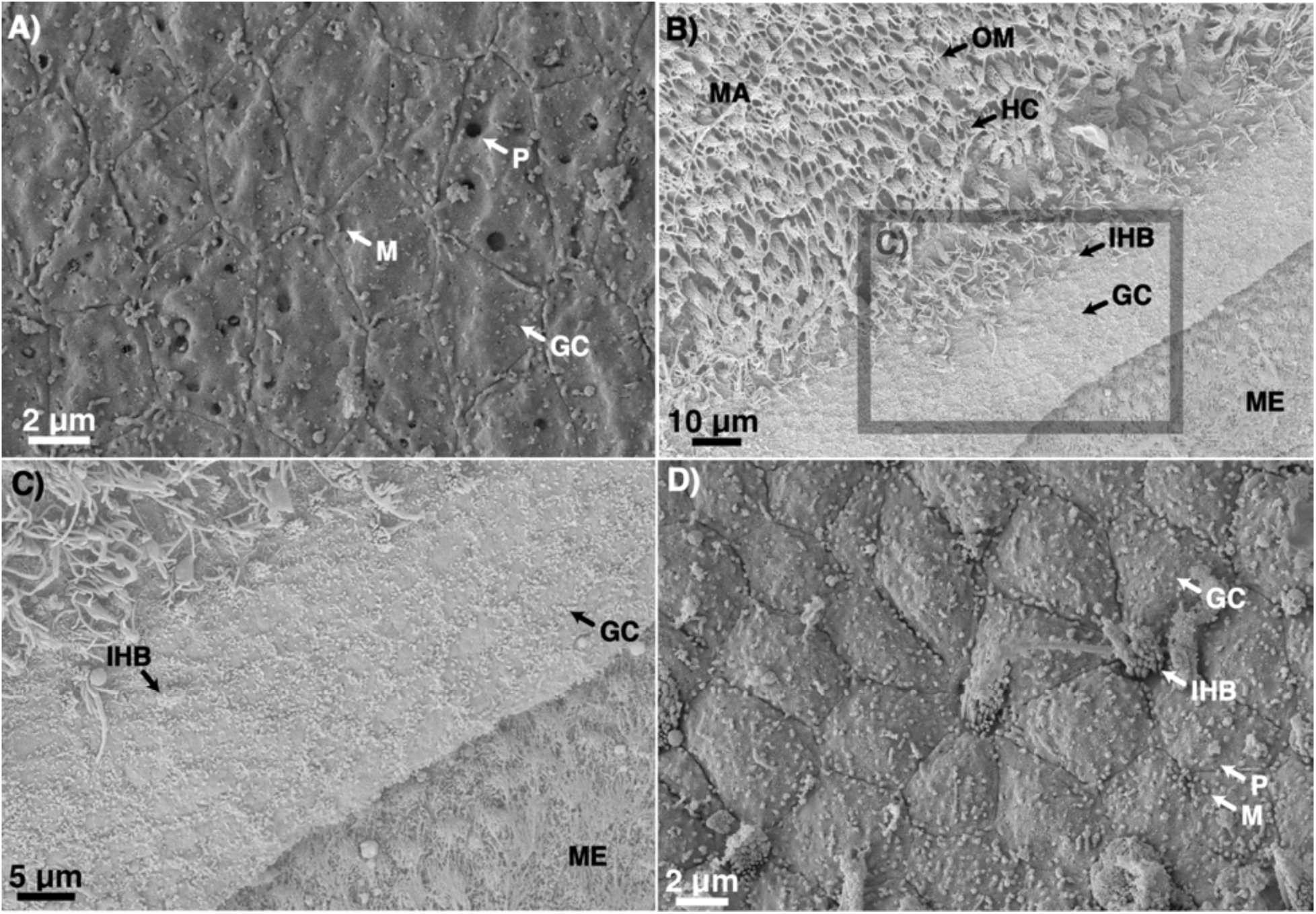
Scanning electron micrographs of inner ear granular cell (GC). A) The apical surface of the GC exhibits numerous microvilli (M) and pores (P). B, C) GCs are located between the macula (MA) and meshwork area (ME), and hair bundles from immature hair cells (IHB) are occasionally observed. D) Like the mature hair cell and supporting cell, IHB are never adjacent to one another and always surrounded by GC. Abbreviation: OM=otolith matrix

The GC nuclei are in the basal part of the cell and form a distinct row in lateral views of three-dimensional confocal reconstructions (Figures 7D, H; 8C, F). The GCs express NKA within their basolateral membrane (Figure 7, 8) and abundant CA within its subapical region (Figure 7A-D). The GCs also contain cytoplasmic VHA and basolateral NKCC at levels similar to the SCs (Figure 7E-H). The most distinguishable feature of the GCs is their high abundance of OMP-1 and Otolin-1 in their apical and sub-apical region (Figure 8). As is the case in HC and SC, the GC’s OMP-1 signal is present at the periphery of the apical membrane (Figure 8B) whereas Otolin-1 signal is in the central part of the apical membrane (Figure 8E). Notably, the OMP-1 signal is very intense in the GCs that are closest to the macula, but they become much dimmer in cells closer to the meshwork area. In contrast, the Otolin-1 signal is uniformly very high across all GCs (Figure 8A, B). Lastly, the expression patterns of CA (Figure 7A-D), NKCC (Figure 7E-H), and OMP-1 resembles that of the otolithic membrane, and this configuration is contiguous and adjoins with that of the SC’s.

**Figure 7:**
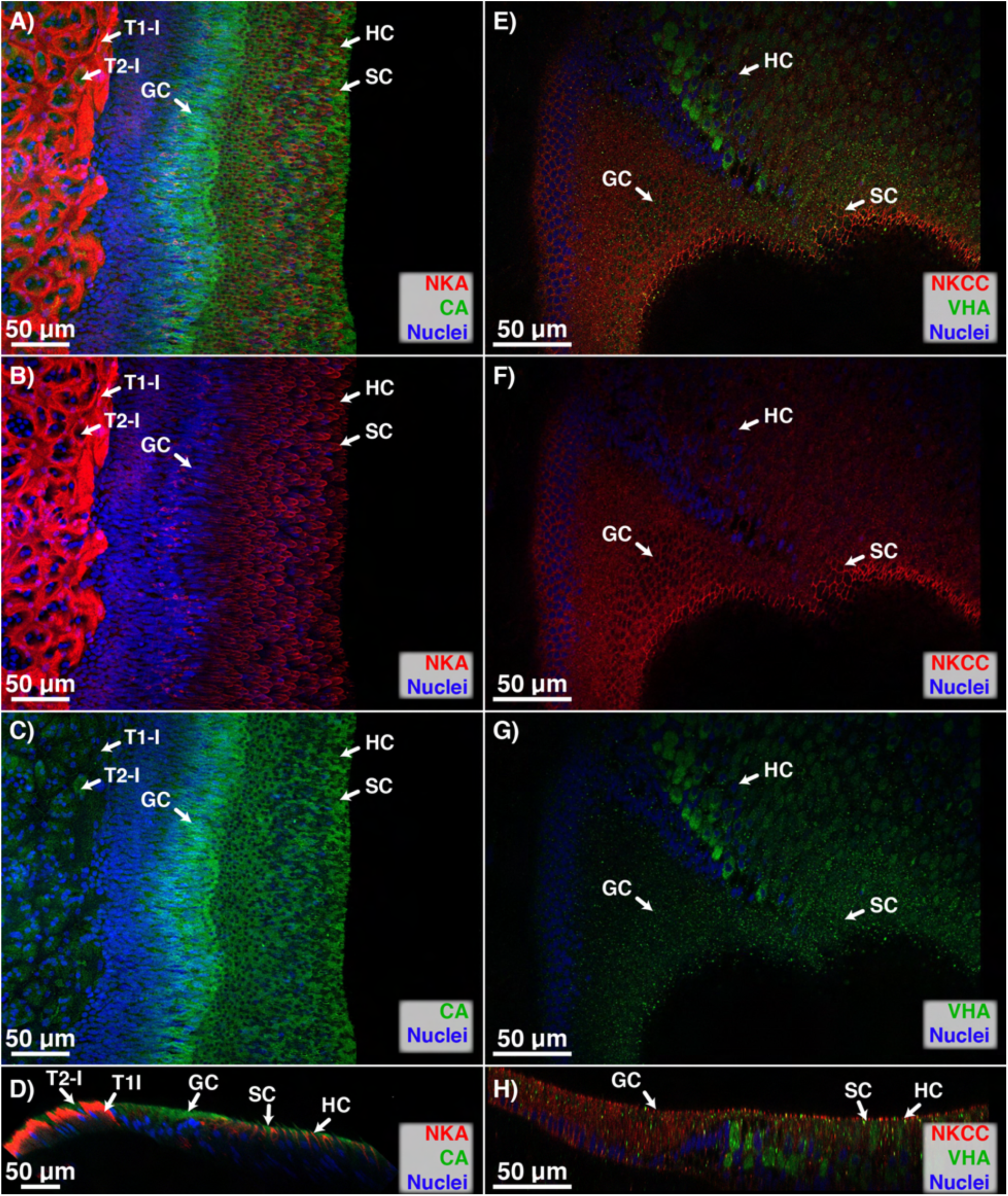
Immunolocalization of NKA, CA, VHA, and NKCC in the granular cell (GC). The GC separates the hair cell (HC) and supporting cell (SC) from the Type-I (T1-I) and Type-II ionocytes(T2-I). A-D) GC expresses basolateral NKA – though not as intensely as the nearby T1-I and HC. A-D) The GC also contains cytoplasmic and basolateral CA and E-F) VHA, and apical and basolateral NKCC. E, F) In particular, NKCC expression resembles the overlying protein matrix, and links to that of the adjacent SC.

**Figure 8:**
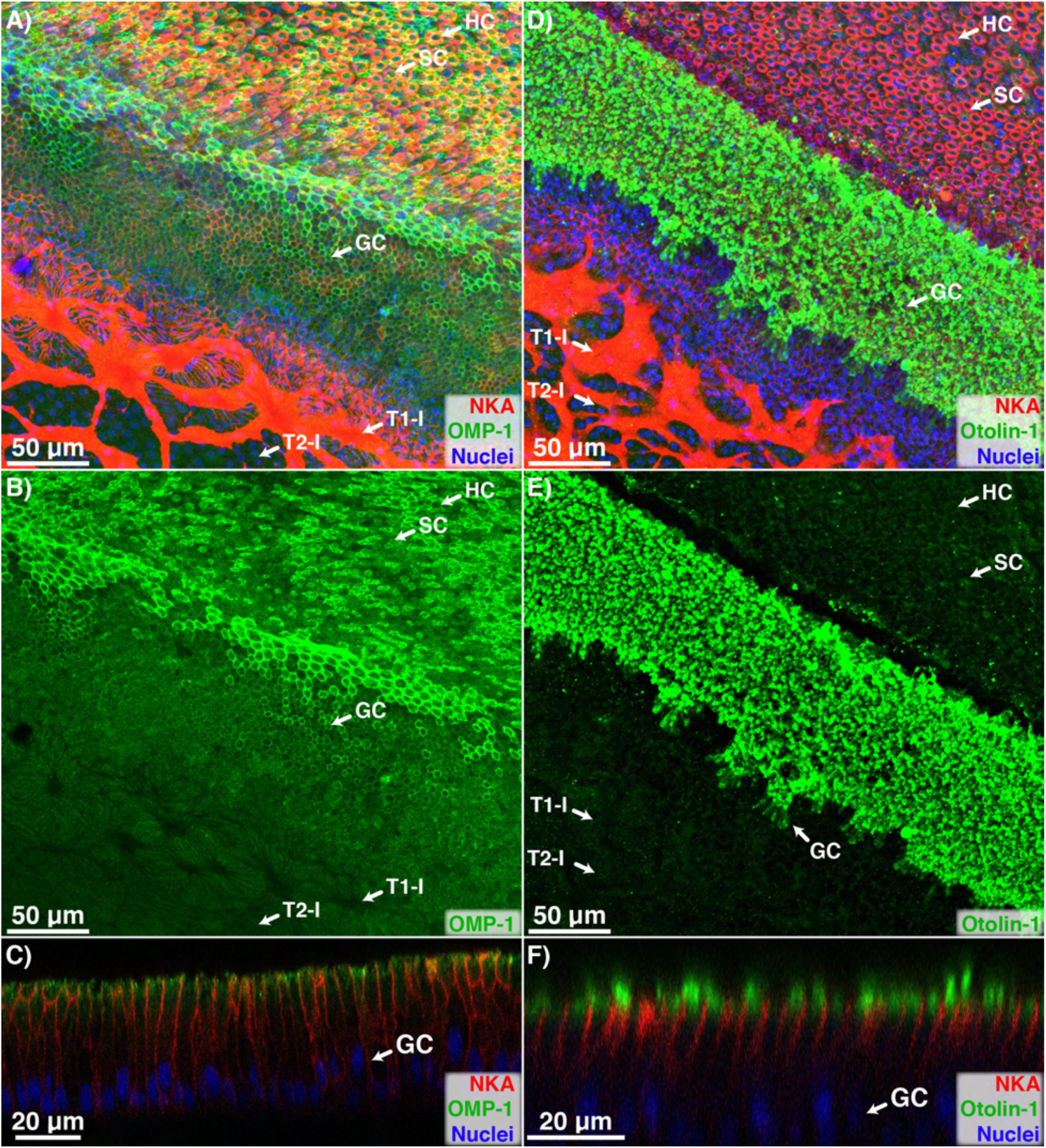
Immunolocalization of NKA, OMP-1, and Otolin-1 in the granular cell (GC). The GC separates the hair cell (HC) and supporting cell (SC) from the Type-I (T1-I) and Type-II ionocytes(T2-I). The GC expresses basolateral NKA, and cytoplasmic and A-C) apical OMP-1 and D-F) Otolin-1. Apical OMP-1 expression resembles the overlying protein matrix, and links to that of the adjacent SC. In contrast, Otolin-1 expression is in the central part of the apical membrane.

### Meshwork and Intermediate Area: Type-I (T1-I) and Type-II (T2-I) Ionocytes

The macula is surrounded by the meshwork area, which is named after the interconnected network of T1-Is observed in past histological studies (Figure 7A, B). SEM imaging outlines a similar meshwork aspect resulting from the contrast in the apical membrane morphology between T1-Is and T2-Is (Figure 9A, B). Higher magnification SEM images (Figure 9C-F) reveal that the T1-Is have more abundant microvilli that the T2-Is, which contains numerous pores (Figure 9G, H). In addition, the microvilli in the T1-Is are significantly shorter (0.568 ± 0.023 μm vs. length: 1.467 ± 0.095 μm; *p* < 0.0001) and wider (0.184 ± 0.006 μm vs. 0.105 ± 0.004 μm; *p* < 0.0001) than those in the T2-Is. As distance from the macula increases, fewer T1-Is were observed, whereas T2-Is become increasingly abundant (Figure 9E-H). This pattern continues until the T1-Is are no longer present, and this signifies the transition from meshwork to intermediate area (Supplemental Figure 3).

**Figure 9:**
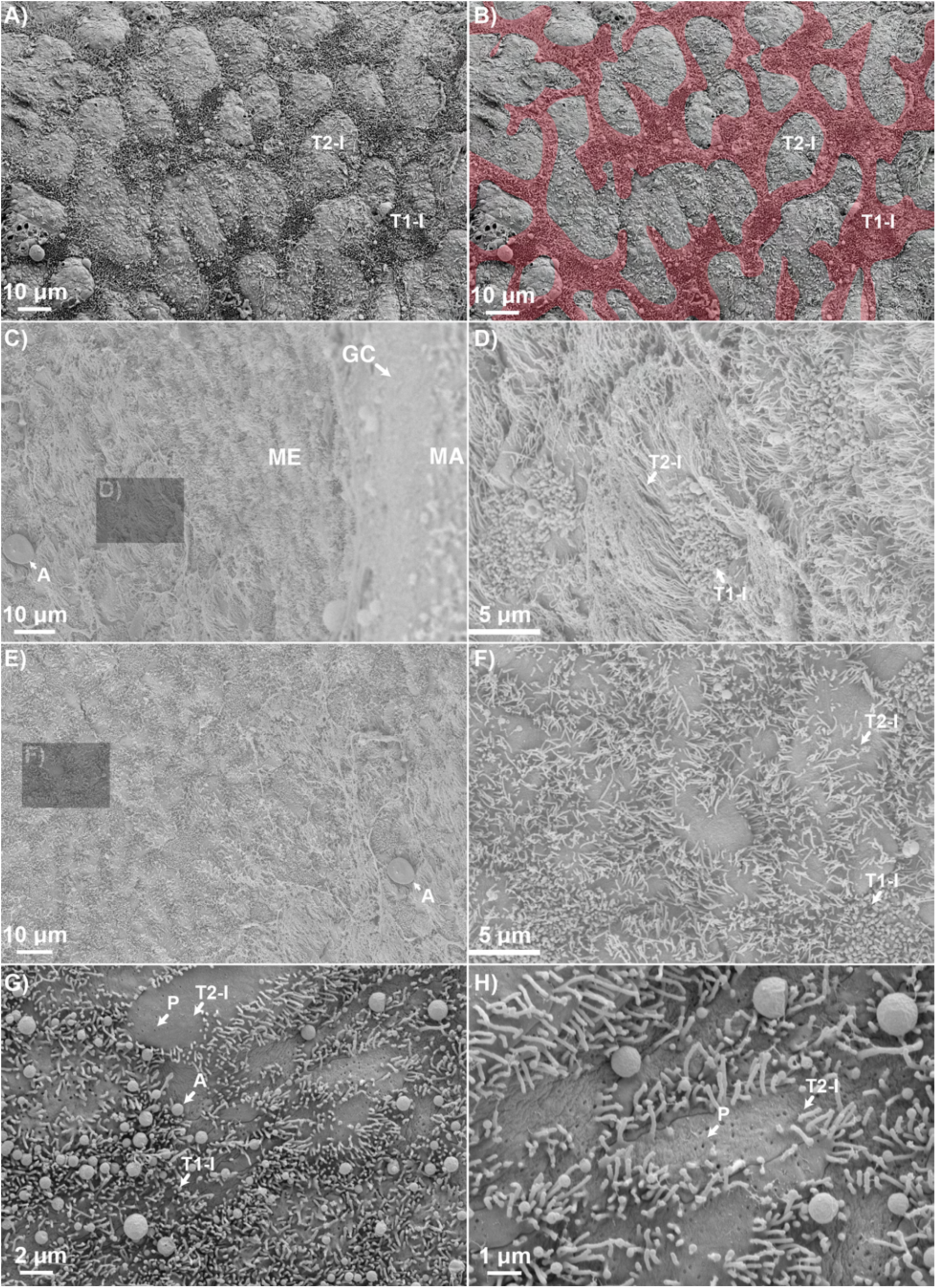
Scanning electron micrographs of inner ear Type-I (T1-I) and T2-I Ionocytes (T2-I). A) The apical surface of the T1-I has more numerous microvilli (M) than the T2-I, and B) matches the interconnected membrane system apparent in basolateral NKA and NKCC immunostaining. C, D) In contrast, the apical surface of the T2-I is relatively more barren, but its microvilli are longer than those of the T1-I. E, F) Microvilli density and length decreases with distance from macula. Please note images C and E are contiguous, and the same artifact (A) is shown as a marker. G, H) Numerous pores (P) can also be observed on the T2-I. Abbreviation: GC=granular cell. ME=meshwork area. MA=macula. A=artifact. OM=otolith matrix

Three-dimensional reconstructions of co-immunostaining combinations indicate that the T1-Is express abundant NKA throughout their basolateral membrane (Figure 7A, B; 8A, D; 10A-D; 11), NKCC in the upper part of the basolateral membrane closer to the apical membrane (Figure 10E-H), and CA (Figure 10A-D) and VHA (Figure 10E-H; 11) throughout the cytoplasm. The T1-Is also have some signal for OMP-1 (Figure 8A, B; 12A-C) and Otolin-1 (Figure 8D, E; 12D-F) in their sub-apical region; however, they were much fainter than in all other inner ear epithelial cell types. Notably, the lack of overlap in immunohistochemical signal between the cytoplasmic CA and VHA and the basolateral NKA (Figure 10A; 11A, D) and NKCC (Figure 10E). Relative to the NKA signal, the NKCC signal in the T1-I appears to be concentrated closer to the apical side of the cell (Figure 10B, F). T1-Is appear to faintly express some OMP-1 (Figure 8A, B; 12A-C) and Otolin-1 (Figure 8D, E; 12D-F) in the sub-apical region of the T1-I.

**Figure 10:**
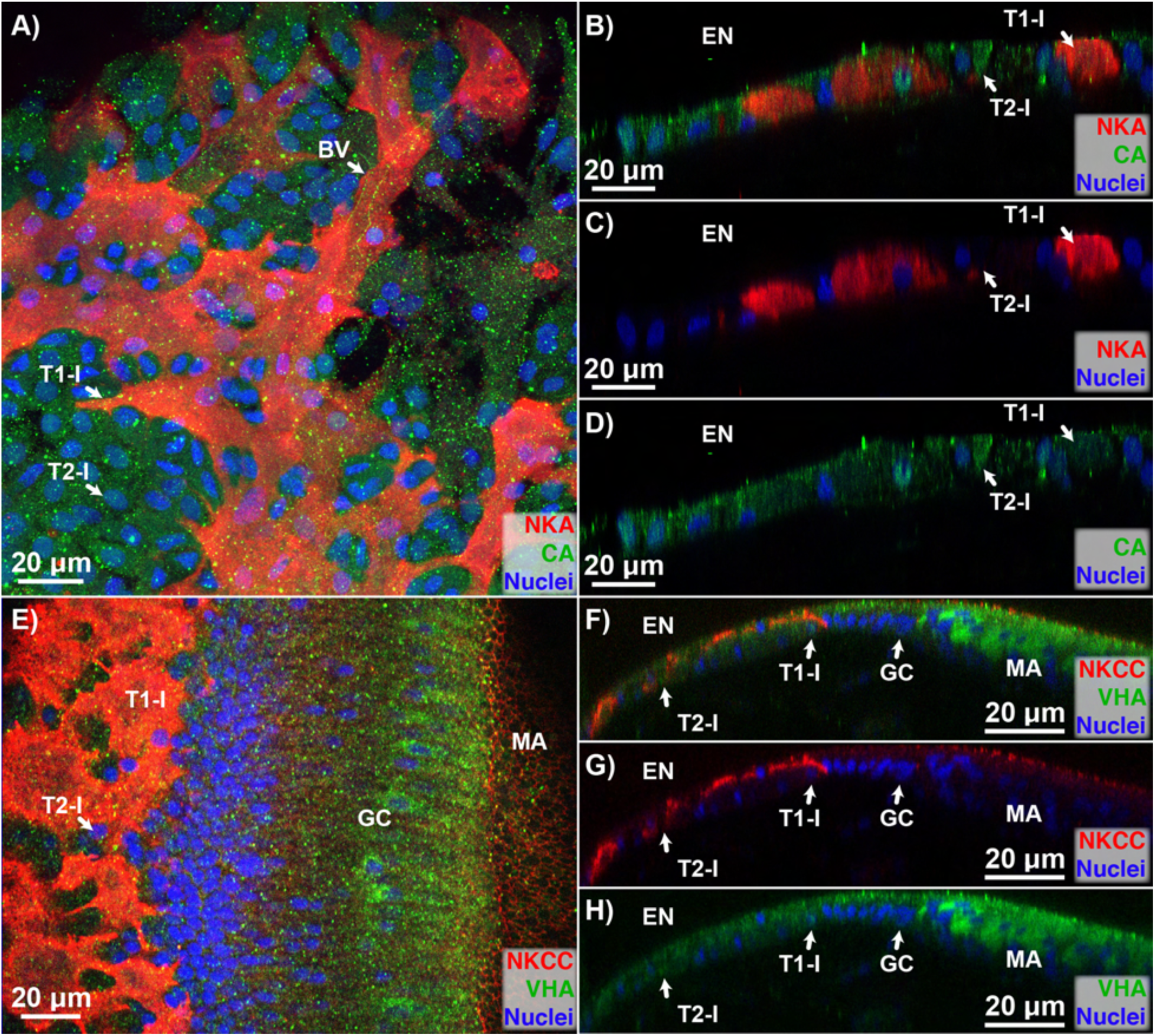
Immunolocalization of NKA, CA, VHA, and NKCC in the Type-I (T1-I) and T2-I Ionocytes (T2-I). The T1-I intensely expresses A-D) basolateral NKA and E-H) NKCC, along with cytoplasmic CA and VHA. In contrast, T2-I does not contain NKA or NKCC, but does express cytoplasmic CA and VHA. Abbreviation: EN=endolymph. MA=macula. GC=granular cell.

**Figure 11:**
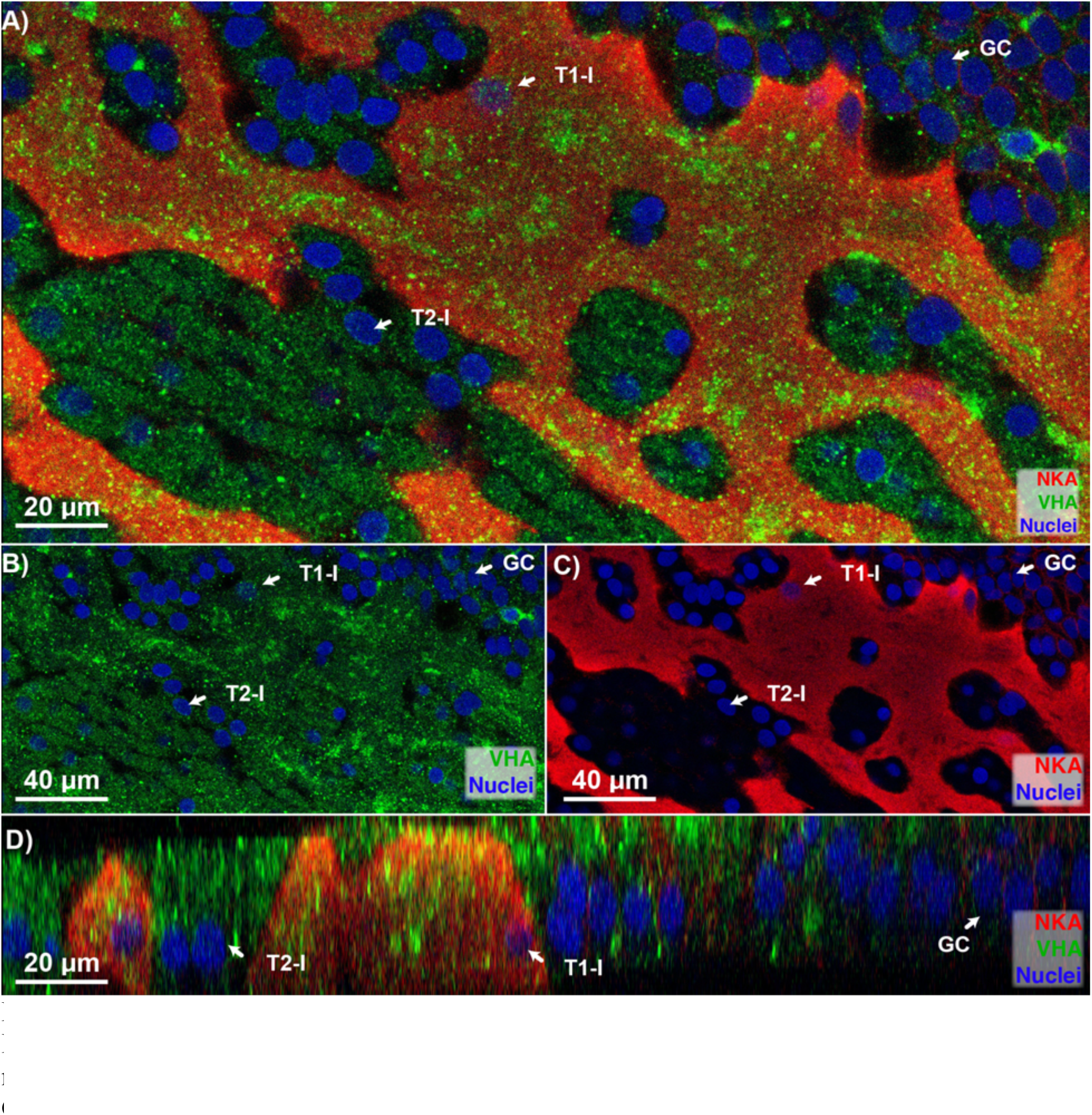
Immunolocalization of NKA and VHA in the Type-I (T1-I) and T2-I Ionocytes (T2-I). A-D) The NKA-rich T1-Is are interconnected with one another, forming a continuous membrane system. In contrast, T2-I do not form a continuous system. Abbreviation: GC=granular cell.

**Figure 12:**
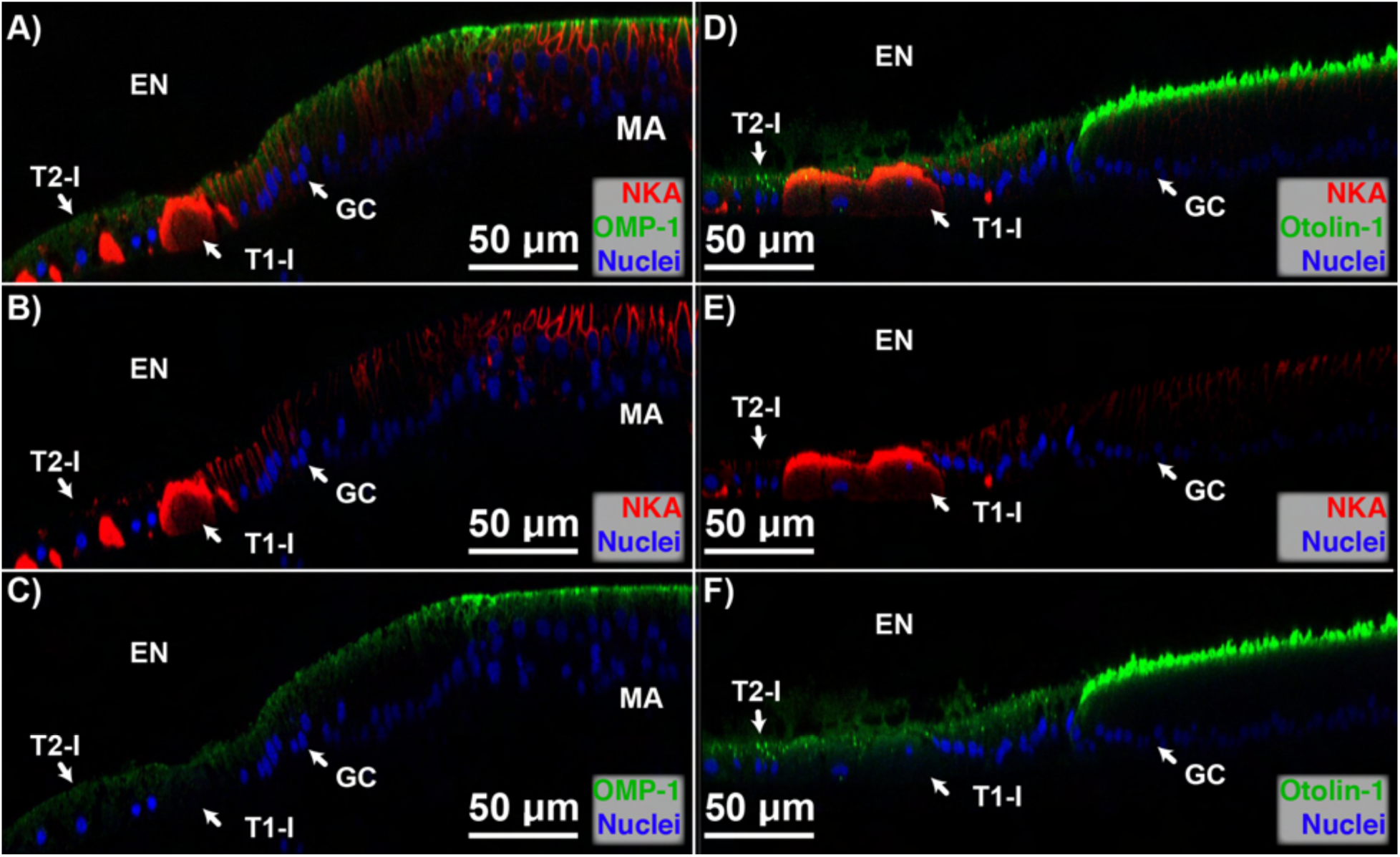
Immunolocalization of NKA, OMP-1, and Otolin-1 in the Type-I (T1-I) and T2-I Ionocytes (T2-I). A-C) OMP-1 is faintly expressed in both the NKA-rich T1-I, and in the T2-I. D-F) Similarly, both T1-I and T2-I appears to express Otolin-1, with parts of it extruding into the endolymph (EN). Abbreviation: MA=macula. GC = granular cell.

In contrast, the T2-Is do not form a continuous membrane system, nor does it express NKA or NKCC (Figures 10, 11). Instead, the T2-Is has abundant CA (Figure 10A-D) and VHA (Figures 10E-H; 11), and faintly express OMP-1 (Figure 12A-C) and Otolin-1 (Figure 12D-F).

### Patches Area: Squamous Cell (SQ)

The patches area is located on the opposite end of the macula and meshwork area, and is entirely comprised of SQs. SEM imaging reveals the border of the SQ contains abundant microvilli, whereas the rest of the cell appears to be relatively barren (Figure 13).

**Figure 13:**
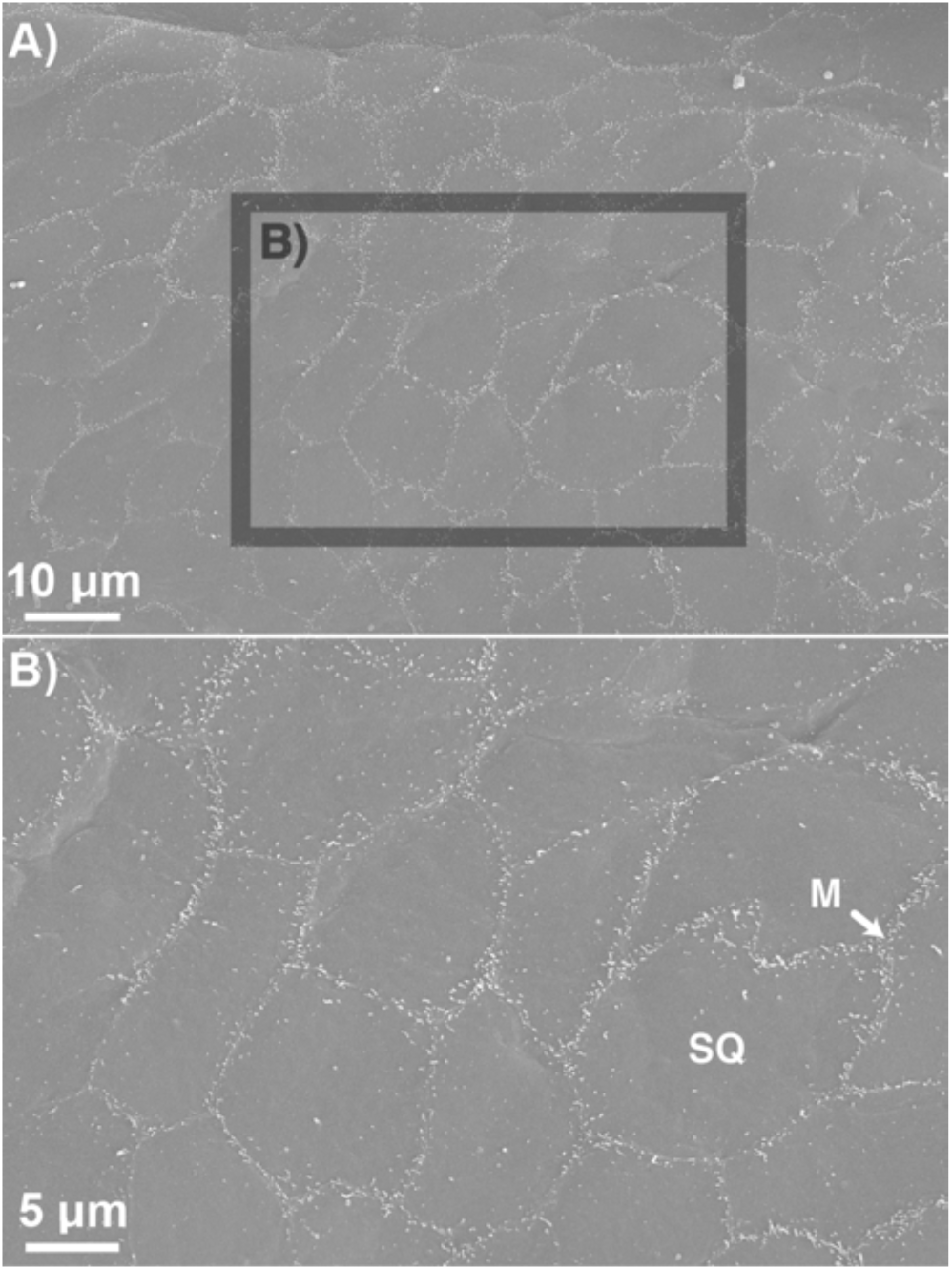
Scanning electron micrographs of inner ear squamous cell (SQ). A) Most of the apical surface on the SQ is relatively barren, B) but its periphery is marked with abundant microvilli (M).

The SQs express abundant NKA and can be mistaken as a T1-I when viewed laterally (Figure 14A, B) or at low-magnification. Instead, the two cell types can be distinguished when viewed apically or at higher magnification settings: the edge of each SQ can be visualized using NKA immunostaining and the position of its nucleus is near the center of the cell (Figure 14C), whereas the borders of the T1-I cannot be defined by NKA and the location of its nucleus do not follow a specific pattern (Figure 11A, D). The SQs also contain abundant cytoplasmic and basolateral CA – with the latter appearing to clearly define the border of its cell membrane (Figure 14C); this immunolocalization pattern sharply contrasts with basolateral NKA and NKCC (Figure 14D) signal along the borders of the cell. VHA is expressed within the cytoplasm of the SQ. Lastly, OMP-1 is expressed along the borders of the SQ (Figure 14E), whereas Otolin-1 is more readily localized in the central portions of the SQ (Figure 14F).

**Figure 14:**
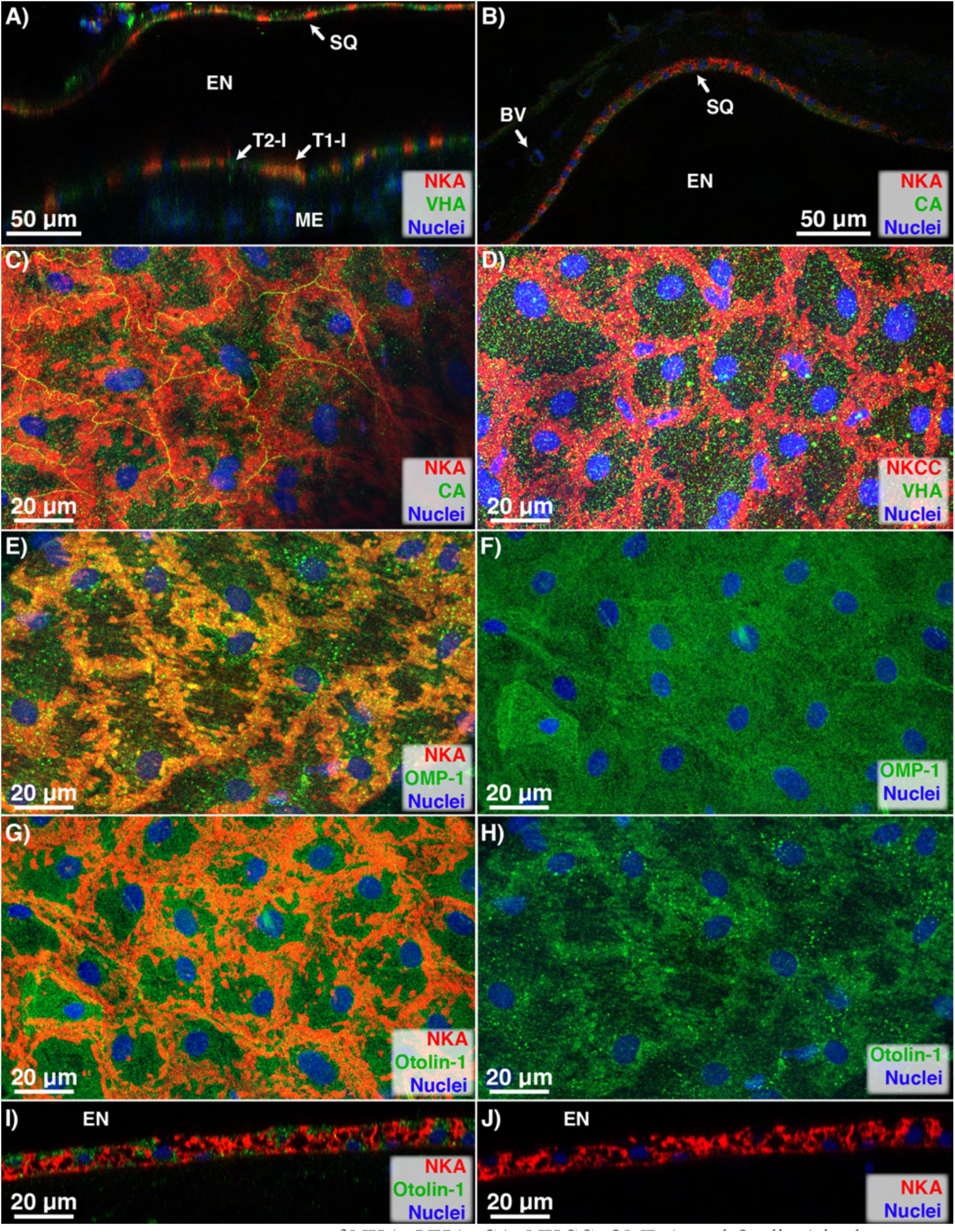
Immunolocalization of NKA, VHA, CA, NKCC, OMP-1, and Otolin-1 in the squamous cell (SQ). A-D) The SQ expresses basolateral NKA and NKCC, and cytoplasmic CA and VHA. E, F) The SQ also expresses OMP-1 near the central part of the cell, but G-J) Otolin-1 at the periphery portions of the cell. Abbreviation: EN=endolymph, T1-I = Type-I Ionocyte, T2-I = Type-II Ionocyte, ME=meshwork area

Altogether, the inner ear cell type-specific intracellular localization and relative protein abundance are summarized in Table 1.

## Discussion

The visualization of the six cell types lining the inner ear endolymph through immunohistochemical and ultrastructural techniques provides the foundational knowledge necessary to understanding the mechanism responsible for inner ear function and otolith biomineralization.

### Hair cell (HC) and supporting cell (SC)

The link between nominal inner ear sensory function and high endolymphatic [K^+^] appears to be evolutionarily conserved across teleost, amphibians, aves, and mammals (63). As the hair bundle is moved by the inertial force of the otolith/otoconia, endolymphatic K^+^ and Ca^2+^ ions rush into the negatively charged HC and rapidly depolarizes the cell (26, 63). During depolarization, the intracellular VHA within the HC may be forming acidic vesicle for neurotransmitter transport for their exocytosis (64), and appears to look similar to the intracellular VHA pattern in nearby neurons. This also explains why nearby SCs have considerably less VHA expression. As the hair bundle reverts back to its original position, intracellular [K^+^] within the HC is removed from the cell via basolateral K^+^ channels (e.g. KCNB1; KCNQ1) (65). Simultaneously, basolateral NKA repolarizes the HC. The removed K^+^ ions are transported away from the HC by the SC, and eventually recycled back into the endolymph. Given that SCs completely envelopes the HC’s basolateral membrane, the role of the SC is likely to assist the HC in repolarization by removing surplus K^+^ as well as resupplying Na^+^ into their interstitial space. The exact mechanism(s) is still being debated, and multiple pathways have been proposed for mammalian and avian models (26, 63). However, the mechanistic pathway for K^+^ recycling in fish likely differ as mammalian Deiters’ cells (analogous to the SCs) have neither NKA nor NKCC while avian Deiters’ cells have both basolateral NKA and NKCC (66), whereas our results suggest teleost SCs do not express NKA but do have apical and basolateral NKCC signal. It is unlikely for a cell to possess both apical and basolateral NKCC. Instead, the apical NKCC signal could instead be a part of the otolithic membrane given that it extends past the soma of the HCs (Figure 4B, C) and is shaped in a pattern resembling that of the otolithic membrane (Figure 2A; 4D, E; 7E, F). Alternatively, the NKCC antibody could simply be mislabeling the intended protein. Future research should focus on identifying the K^+^ channel within teleost HC and SC, and verifying the presence of NKCC in the SC.

In addition to NKCC, SCs also have CA and OMP-1 signals matching the otolithic membrane pattern observed in the SEM. This suggests the SCs play a critical role in secreting the otolithic membrane that protects the adjacent HC from being crushed by the overlying otolith, as well as anchoring the matrix structure to the macula (20, 28, 67). This otolithic membrane must expand outwards to support the perennial growth of the otolith, yet remain porous to allow endolymph K^+^ ions to reach the HCs at the center of the macula. Some SC also appears to express Otolin-1 in the middle of its apical membrane, which suggests it may be contributing to the biomineralization of the otolith near the central region of the macula (30–32). Interestingly, previous Otolin-1 immunostaining on the chum salmon (*Oncorhychus keta*) and the rainbow trout (*Oncorhynchus mykiss*) were immunonegative (31, 43). Perhaps the prevalence of Otolin-1 within the inner ear is linked to the relative otolith size: rockfishes (*Sebastes spp.*) are notorious for having large otoliths, whereas salmonids generally have relatively smaller otoliths.

### Granular Cell (GC)

Similar to previous studies (31, 43), Otolin-1 presence is significantly greater within the GC than in any other inner ear cell type. This suggests the GC is not only critical to otolith biomineralization, but also necessary for proper CaCO_3_ arrangement: calcite has a rhombohedral crystal structure and is formed in the presence of exclusively Otolin-1, whereas aragonite has an orthorhombic crystal structure formation that requires the presence of both Otolin-1 and otolith matrix molecule 64 (OMM-64) during *in vitro* trials (30, 68). The abundant presence of Otolin-1 in the GC around the macula also matches the natural biomineralization patterns: the proximal side (facing the macula) is known to grow faster and have wider increments than the distal side (facing the patches area) of the otolith (40).

In contrast, OMP-1 expression is not uniform. OMP-1 is more abundantly expressed in the GCs closest to the macula, and this spatial pattern suggests their role in supporting and accommodating the growing otolith. As the otolith grows, so must new HC and SC be synthesized and expand outward – thereby displacing the adjacent GC. In this study, incipient HCs are observed in the midst of GCs at a significant distance away from the rest of the macula (Figure 6). As such, do GCs gradually morph into SCs? This may also explain why the SC and GC share similar CA, VHA, and basolateral NKCC expression (both in intracellular localization and relative abundance). Other proteins critical to otolith biomineralization that the GCs may be secreting include *starmaker* (69, 70) and *OMM-64* (68), though species-specific variation exists [c.f. *Oreochromis mossambicus* lacks OMM-64 and have calcitic otoliths (71)]. There is much that is not known about GC, and further research is necessary to confirm whether GCs are displaced or transforms into SCs as the otolith enlarges, and to verify whether they secrete other otolith-relevant proteins.

### Type-I (T1-I) and Type-II Ionocytes (T2-I)

The primary role of the T1-I has long been proposed to be responsible for maintaining the ionic and acid-base balance of the inner ear endolymph given its ample expression of mitochondria (18, 38), NKA (24, 37, 54), NKCC (37, 59), and the potassium channel KCNQ1 (59). In fact, the proposed mechanism for K^+^ secretion within the inner ear appears to be evolutionary conserved (63, 66): the teleost T1-I (37) seems to function very similarly to their analogous counterparts, the mammalian marginal cells (72, 73) and the avian vestibular dark cells (74). However, and unlike their mammalian and avian counterparts, the teleost inner ear is unique in that their otoliths biomineralizes continuously throughout the fish’s life.

In contrast, the role of T2-I does not express NKA nor NKCC, and it likely doesn’t play a role in K^+^ secretion that the T1-I specializes in. Given this, the T2-I may simply play a role in maintaining the ionic gradients generated by the T1-I. This likely requires the expression of tight junctions via a claudin-like protein to prevent ion leakage and to maintain the endolymphatic ionic K^+^ gradient similar to the inner ears of mice (18, 75, 76).

Continuous H^+^ removal is necessary to promote perennial otolith biomineralization, and both T1-I and T2-I express CA and VHA, two proteins often involved in H^+^/HCO_3_^-^ transport. The role of H^+^ removal is often linked to VHA since it has been shown to translocate from the cytoplasm to the cell membrane during acid-base disturbance (77–79). Even so, VHA does not appear to play a role in mitigating an environmentally-relevant acid-base disturbance (24, 37). Instead, VHA may be forming acidic vesicles to sequester H^+^ within the T1-I in order to promote the formation of amorphous CaCO_3_ similar to the calcifying primary mesenchyme cells within the sea urchin (80). This may explain the presence of CA, which would catalyze local CO_2_ into H^+^ and HCO_3_^-^ to the formation of amorphous CaCO_3_ within the T1-I. The use of acidic vesicles to promote amorphous CaCO_3_ has long been proposed for teleost otolith biomineralization (81–83), and there are evidence of this mechanism in calcifying corals (84–86). Further research is necessary to confirm whether the VHA expression within the T1-I are indeed intracellular vesicles involved in sequestering and removing H^+^ to promote biomineralization.

The utilization of CA and VHA to generate H^+^ necessitates the concurrent expression of a HCO_3_^-^ transporter, and like other models pendrin is the likely culprit. Pendrin (*slc26a4*) is an anion exchanger that has been demonstrated to be critical to hearing in mammalian models, and knockout mice have more acidic endolymph pH and higher Ca^2+^ (87, 88). This suggests an anion exchanger could be present on the apical membrane to transport HCO_3_^-^ into the endolymph to buffer H^+^ accumulation, or on the basolateral membrane to amass HCO_3_^-^ in the endolymph to promote biomineralization. However, the localization of an anion exchanger within fish remains to be discovered, and further research is necessary to determine its intracellular expression within the inner ear cell type(s).

Both T1-I and T2-I appears to faintly express OMP-1 and/or Otolin-1 near the sub-apical and apical portions of the cells, and the two proteins are likely involved with intracellular synthesis of amorphous CaCO_3_ as proposed in sea urchins (80). OMP-1 and Otolin-1 staining appears to be extruded from the T1-I and T2-I into the endolymph (Figure 12) in a manner similar to past TEM observations (89). Furthermore, SEM analysis revealed numerous pores suggesting exocytosis. Although the initial reporting within the rainbow trout did not immunolocalize OMP-1 or Otolin-1 in T1-I and T2-I (43), this could be due to differences in the sensitivity of the microscope and technique. Even so, additional studies are needed to determine whether OMP-1, Otolin-1, or another similar matrix protein is present within the T1-I and T2-I, and whether it is involved in synthesizing amorphous CaCO_3_.

### Squamous Cell (SQ)

SQ is the only cell type facing the distal portion of the otolith, and its roles include ion-transport (42) and otolithic protein secretion (31). The co-localized expression of NKA and NKCC and the extended microvilli along the perimeter of the cell resembles that of the T1-I, and thus SQ is likely also generating an electrochemical gradient for ion-transport. Given the basolateral NKA and NKCC and the endolymph’s elevated [K^+^], the potassium channel KCNQ1 is likely expressed on the apical membrane to facilitate K^+^ secretion (59). Unlike the macula and meshwork area, the patches area is known to be particularly thin and have relatively few capillaries (18). Thus, the SQs could be sourcing their ions from the perilymph (rather than from blood capillaries). Yet unlike the T1-I, the perimeter of the SQ also expresses CA. Moreover, the presence of Otolin-1 and OMP-1 have been observed (43). While the exact mechanism remains unknown, SQs are likely integral in maintaining the nominal endolymph chemistry and generating the necessary substrates for otolith biomineralization on the distal side of the otolith (40).

### Proximal-Distal Gradient and Growth-Influenced Alteration

Altogether, the spatial position, cell specialization, and endolymph chemistry align with past observations on proximal-distal ionic gradient (19, 40) and relative number of amorphous CaCO_3_ (34, 35). The network of mitochondrion-rich T1-I and the abundant GC located on the proximal side of the inner ear are likely able to significantly transport greater ions and synthesize more proteins compared to the SQ on the distal side of the inner ear. To elaborate, the relatively lower [K^+^] in the proximal endolymph (19, 40) likely stems from its utilization by the HCs and removal by SCs. Next, the relatively lower pH in the proximal endolymph (19) suggests greater biomineralization (which produces H^+^) and cellular respiration (which produces CO2 that hydrates into HCO_3_^-^ and H^+^). Furthermore, the relatively greater protein concentration within the proximal endolymph (40) suggests the outsized production by the GC. Finally, there are more amorphous CaCO_3_ both in the endolymph and bound to the edge of the otolith (34, 35), which ultimately leads to greater biomineralization on the proximal face of the otolith (33). Conversely, the distal endolymph is bordered by the SQ, which has significantly less mitochondria (38) and NKA, and must fulfill the role of multiple cell types. As such, the distal endolymph has relatively higher [K^+^] and pH (19, 40), lower protein concentration (40), less amorphous CaCO_3_ (34, 35), and slower otolith biomineralization (33).

The otolith grows continuously throughout the fish’s life, and thus the inner ear organ must also remodel itself to meet the demand. As the otolith expands, the need for additional matrix protein to protect the underlying HC is necessary. This means the HC and SC in the macula must expand outward, and this study provides evidence that incipient HC is synthesized in the area occupied by GC. However, it is unclear how new SC are formed: do GCs become SCs, or would new SCs be synthesized? Nevertheless, as the GC becomes displaced, new GCs must be created in the area that the T1-I and T2-I are located. This results in the displacement of T1-I towards the Intermediate Area, and could explain why the T1-I at the edge of the macula have a “scrunched up” appearance.

The capacity for the inner ear to remodel itself suggests it could also balance and control the demands for ion-transport and otolith biomineralization. Firstly, the presence of extended microvilli across most of the described cell types suggest a degree of control of activities: increased ion-transport could be generated with greater extension, whereas slowing of this process is possible with microvilli retraction. Moreover, as the inner ear grows, there is a potential for adjusting the number of cell types: the synthesis of more GC could indicate the need to boost otolith growth, whereas increasing T1-I may suggest the need for additional [K^+^] to meet HC demands. Altogether, this suggests the inner ear is capable of remodeling, and could potentially alter its protein abundance, apical morphology, and adjust the number of cell types in response to chronic stressors much like the gill ionocytes to various environmental stressors (c.f. (90–92). As a result, it remains possible that the inner ear’s long-term response to future ocean acidification, warming, and hypoxia may not be predictable using short-term studies. Future research focused on understanding the dynamics of inner ear control over otolith biomineralization and the interplay between cell types could potentially explain why otolith size varies across species.

### Implications for Otolith-Reliant Tools under Future Climate Change Scenarios

This study showcases the immense complexities of the inner ear, highlights numerous questions that remains unanswered, and demonstrates the urgent need to understand the mechanism underpinning otolith biomineralization. Already, otolith biomineralization has already been shown to be impacted by future climate change scenarios including ocean warming (47, 48), acidification (45, 46), and hypoxia (49–51). Moreover, studies on otolith responses to multiple stressors are scarce: ocean acidification and warming impacts are synergistic and led to significantly greater otolith size (48), but the effects on ocean acidification and hypoxia remain undocumented in fish. However, in embryonic market squid (*Doryteuthis opalescens*) statolith (which is analogous to fish otolith), combined ocean acidification and hypoxia exposure resulted in significantly smaller statolith (93). In summary, future climate change threatens to upend the decades of work spent to build the otolith-reliant toolkit that fishery managers, ecologists, and conservation biologists rely upon. This body of work showcases the need to consider inner ear cell responses in order to more accurately anticipate and predict otolith biomineralization patterns under future climate change scenarios.

Emerging technology such as using Fourier Transformed Near-Infrared Spectroscopy (FT-NIRS) to fish age by estimating its protein absorbance (94–97) is also at risk. Although FT-NIRS has the potential to greatly increase cost-efficiency and reduce human error, climate change could still introduce unanticipated errors to the FT-NIRS by altering the ratio of otolithic proteins (e.g. OMP-1, Otolin-1) and CaCO_3_ as biomineralization rate increases due to ocean warming and acidification, or as biomineralization is stymied by hypoxia. Moreover, the remodeling of the inner ear (similar to that of fish gills) could also otolith biomineralization patterns, elemental incorporation, and protein production/abundance. As such, there remains an immense need for long-term studies that examines the future impacts of ocean warming, acidification, and hypoxia on the otolith metrics (e.g. annulus, elemental and isotopic signature, protein abundance) that fishery managers, ecologists, and conservation biologists rely upon.

## Supplemental Material

External link to supplemental data: https://github.com/gkwan09/SupplementalData

## Acknowledgements

We are grateful to Dr. Emi Murayama (Institut Pasteur, Paris, France) for her donation of the OMP-1 and Otolin-1 antibodies. We thank Dr. Uri Manor for the use of the scanning electron microscope at Salk Institute. We also thank Phil Zerofski (UCSD-SIO) for his help in collecting the juvenile splitnose rockfish, and thank Taylor Smith (UCSD-SIO), Shane Finnerty (UCSD-SIO), and Gabriel Lopez (UCSD-SIO) for their assistance in animal care and rearing. GTK was funded by the NSF Postdoctoral Research in Biology National Science Foundation (NSF) Postdoctoral Research Fellowship in Biology (Award #1907334) and by the Delta Stewardship Council Delta Science Program under Grant No. (21045). The contents of this material do not necessarily reflect the views and policies of the Delta Stewardship Council, nor does mention of trade names or commercial products constitute endorsement or recommendation for use.

## Abbreviations

CaCO_3_: Calcium Carbonate
NKA: Na^+^/K^+^-ATPase
CA: Carbonic Anhydrase
VHA: Vacuolar H^+^-ATPase
NKCC: Na^+^-K^+^-2Cl^-^-Co-Transporter
OMP-1: Otolith Matrix Protein-1
HC: hair cell
SC: supporting cell
GC: granular cell
T1-I: Type-I ionocyte
T2-I: Type-II ionocyte
SQ: squamous cell

## Supplemental Information

**Supplemental Figure 1:**
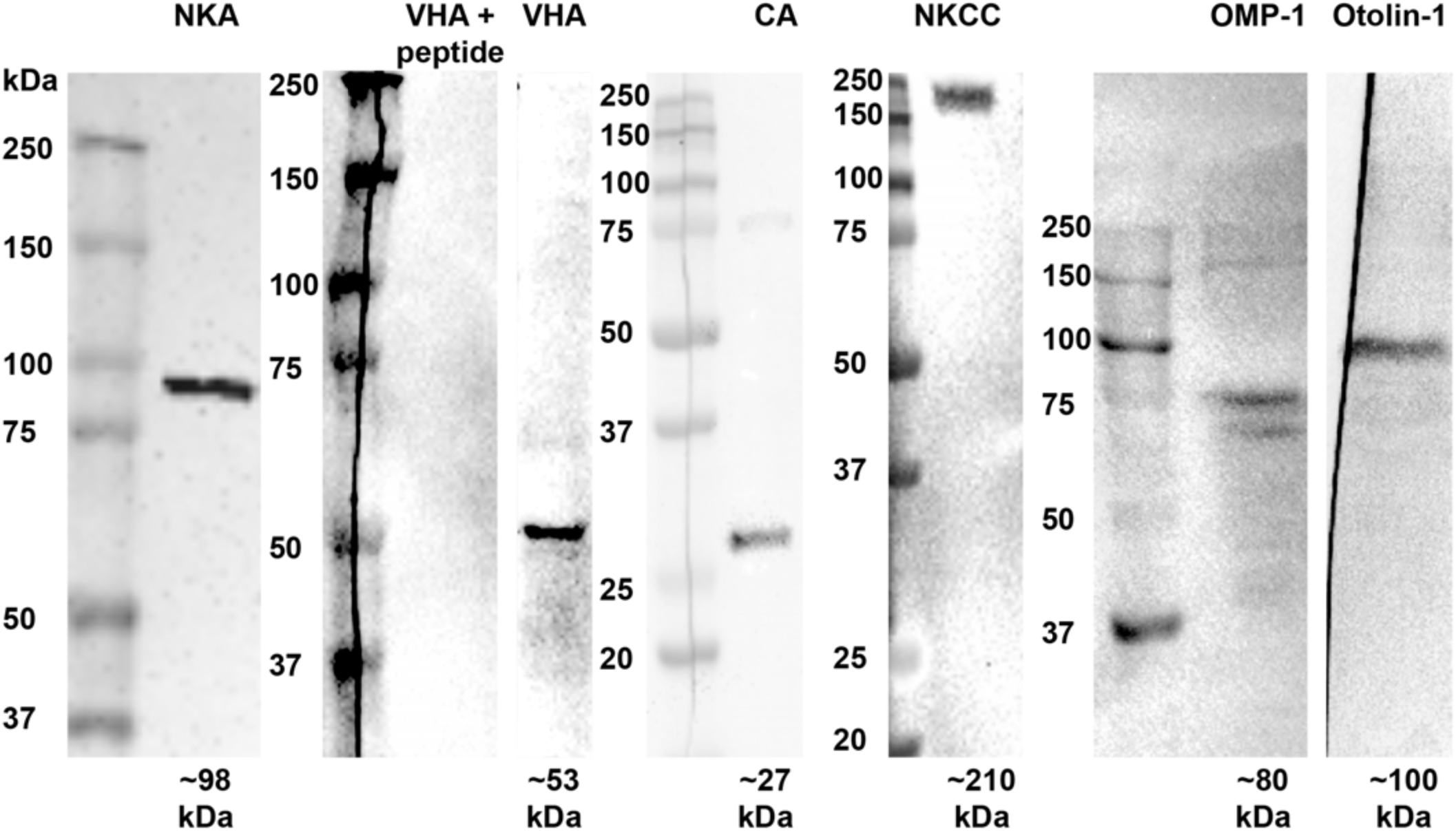
Western blot analysis of splitnose rockfish inner ear homogenates. Antibodies against Na^+^/K^+^-ATPase (mNKA), V-type H^+^ ATPase (VHA), carbonic anhydrase (CA), Na^+^-K^+^-Cl^−^-co-transporter (NKCC), otolith matrix protein 1 (OMP-1), and otolin-1 reveal bands matching the predicted size of respective proteins. Molecular marker is shown on the left of each respective blot

**Supplemental Figure 2:**
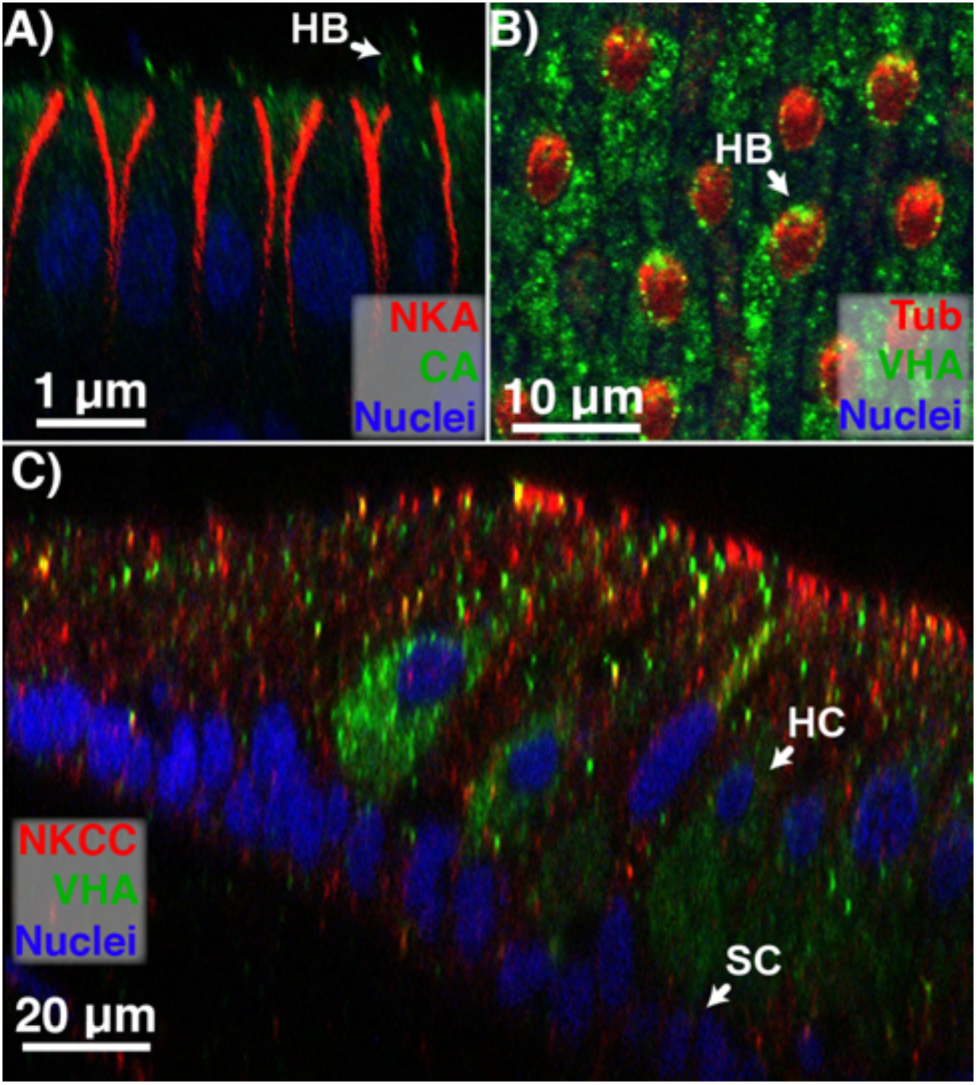
Immunolocalization of NKA, CA, VHA, Tubulin, and NKCC in hair cell (HC) and supporting cell (SC). The NKA-rich HC expresses both A) CA and B) VHA in its hair bundles (HB). The nuclei of the HC and SC forms two different parallel lines. C) When viewed laterally, the two rows of nuclei can be used to distinguish between HC and SC.

**Supplemental Figure 3:**
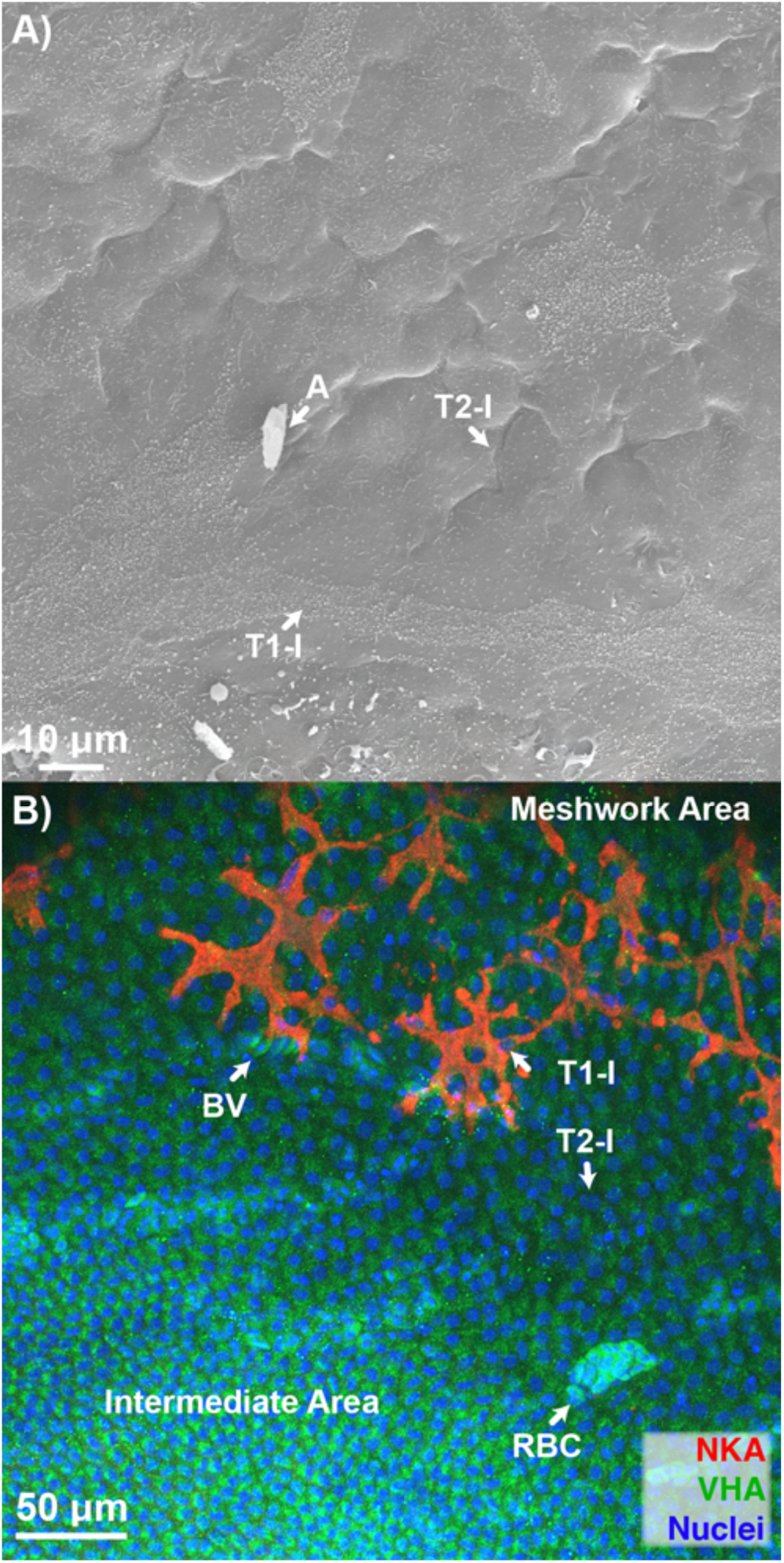
Scanning electron micrograph and immunolocalization of NKA and VHA of Type-I (T1-I) and T2-I Ionocytes (T2-I) at the intermediate area. A) T1-I near the intermediate area is much sparser relative to those near the macula. B) Similarly, the apical surface and microvilli of T2-I in the intermediate area are relatively barren and receded, respectively.

## References

1. Campana SE. Otolith science entering the 21st century. Mar Freshw Res 56: 485–495, 2005.

2. Trofimova T, Alexandroff SJ, Mette MJ, Tray E, Butler PG, Campana SE, Harper EM, Johnson ALA, Morrongiello JR, Peharda M, Schöne BR, Andersson C, Andrus CFT, Black BA, Burchell M, Carroll ML, DeLong KL, Gillanders BM, Grønkjær P, Killam D, Prendergast AL, Reynolds DJ, Scourse JD, Shirai K, Thébault J, Trueman C, de Winter N. Fundamental questions and applications of sclerochronology: Community-defined research priorities. Estuar Coast Shelf Sci 245, 2020. doi: 10.1016/j.ecss.2020.106977.

3. Begg GA, Campana SE, Fowler AJ, Suthers IM. Otolith research and application: current directions in innovation and implementation. Mar Freshw Res 56: 477, 2005. doi: 10.1071/MF05111.

4. Campana SE, Neilson JD. Microstructure of Fish Otoliths. Can J Fish Aquat Sci 42: 1014–1032, 1985.

5. Pannella G. Fish Otoliths : Daily Growth Layers and Periodical Patterns. Science (80-) 173: 1124–1127, 1971.

6. Methot RD. Prioritizing Fish Stock Assessments. NOAA Tech Memo NMFS-F/SPO*-*152: 31 p., 2015.

7. Botsford LW, Holland MD, Samhouri JF, White JW, Hastings A. Importance of age structure in models of the response of upper trophic levels to fishing and climate change. ICES J Mar Sci 68: 1270–1283, 2011. doi: 10.1093/icesjms/fsr042.

8. Alewijnse S, Stowasser G, Saunders R, Belcher A, Crimmen O, Cooper N, Trueman C. Otolith-derived field metabolic rates of myctophids (family Myctophidae) from the Scotia Sea (Southern Ocean). Mar Ecol Prog Ser 675: 113–131, 2021. doi: 10.3354/meps13827.

9. Chung MT, Trueman CN, Godiksen JA, Holmstrup ME, Grønkjær P. Field metabolic rates of teleost fishes are recorded in otolith carbonate. Commun Biol 2: 1–10, 2019. doi: 10.1038/s42003-018-0266-5.

10. Willmes M, Lewis LS, Davis BE, Loiselle L, James HF, Denny C, Baxter R, Conrad JL, Fangue NA, Hung TC, Armstrong RA, Williams IS, Holden P, Hobbs JA. Calibrating temperature reconstructions from fish otolith oxygen isotope analysis for California’s critically endangered Delta Smelt. Rapid Commun Mass Spectrom 33: 1207–1220, 2019. doi: 10.1002/rcm.8464.

11. Janak JM, Linley TJ, Harnish RA, Shen SD. Partitioning maternal and exogenous diet contributions to otolith87sr/86sr in kokanee salmon (Oncorhynchus nerka). Can J Fish Aquat Sci 78: 1146–1157, 2021. doi: 10.1139/cjfas-2020-0242.

12. Limburg KE, Walther BD, Lu Z, Jackman G, Mohan J, Walther Y, Nissling A, Weber PK, Schmitt AK. In search of the dead zone: Use of otoliths for tracking fish exposure to hypoxia. J Mar Syst 141: 167–178, 2015. doi: 10.1016/j.jmarsys.2014.02.014.

13. Shirai K, Koyama F, Murakami-Sugihara N, Nanjo K, Higuchi T, Kohno H, Watanabe Y, Okamoto K, Sano M. Reconstruction of the salinity history associated with movements of mangrove fishes using otolith oxygen isotopic analysis. Mar Ecol Prog Ser 593: 127–139, 2018. doi: 10.3354/meps12514.

14. Swearer SE, Caselle JE, Lea DW, Warner RR. Larval retention and recruitment in an island population of a coral-reef fish. Nature 402: 799–802, 1999. doi: 10.1038/45533.

15. Campana S. Chemistry and composition of fish otoliths: pathways, mechanisms and applications. Mar Ecol Prog Ser 188: 263–297, 1999. doi: 10.3354/meps188263.

16. Green BS, Mapstone BD, Carlos G, Begg GA. Tropical Fish Otoliths: Information for Assessment, Management, and Ecology. Springer, 2009.

17. Hunter E, Laptikhovsky V V., Hollyman PR. Innovative use of sclerochronology in marine resource management. Mar Ecol Prog Ser 598: 155–158, 2018. doi: 10.3354/meps12664.

18. Pisam M, Payan P, LeMoal C, Edeyer A, Boeuf G, Mayer-Gostan N. Ultrastructural study of the saccular epithelium of the inner ear of two teleosts, Oncorhynchus mykiss and Psetta maxima. Cell Tissue Res 294: 261–270, 1998. doi: 10.1007/s004410051176.

19. Payan P, De Pontual H, Bœuf G, Mayer-Gostan N. Endolymph chemistry and otolith growth in fish. Comptes Rendus Palevol 3: 535–547, 2004. doi: 10.1016/j.crpv.2004.07.013.

20. Steinacker A, Menton DN, Romero A. Toadfish saccular hair cell bundle has a preferred orientation in the otolithic membrane. Hear Res 48: 145–149, 1990. doi: 10.1016/0378-5955(90)90204-3.

21. Popper AN. Ultrastructure of the sacculus and lagena in a moray eel (Gymnothorax sp.). J Morphol 161: 241–256, 1979. doi: 10.1002/jmor.1051610302.

22. Dunkelberger DG, Dean JM, Watabe N. The ultrastructure of the otolithic membrane and otolith in the juvenile mummichog, Fundulus heteroclitus. J Morphol 163: 367–377, 1980. doi: 10.1002/jmor.1051630309.

23. Shiao JC, Lin LY, Horng JL, Hwang PP, Kaneko T. How can teleostean inner ear hair cells maintain the proper association with the accreting otolith? J Comp Neurol 488: 331–341, 2005. doi: 10.1002/cne.20578.

24. Kwan GT, Tresguerres M. Elucidating the acid-base mechanisms underlying otolith overgrowth in fish exposed to ocean acidification. Sci Total Environ 823: 153690, 2022. doi: 10.1016/j.scitotenv.2022.153690.

25. Bang PI, Sewell WF, Malicki JJ. Morphology and cell type heterogeneities of the inner ear epithelia in adult and juvenile zebrafish (Danio rerio). J Comp Neurol 438: 173–190, 2001. doi: 10.1002/cne.1308.

26. Zdebik AA, Wangemann P, Jentsch TJ. Potassium ion movement in the inner ear: Insights from genetic disease and mouse models. Physiology 24: 307–316, 2009. doi: 10.1152/physiol.00018.2009.

27. Monroe JD, Rajadinakaran G, Smith ME. Sensory hair cell death and regeneration in fishes. Front Cell Neurosci 9: 1–18, 2015. doi: 10.3389/fncel.2015.00131.

28. Kachar B, Parakkal M, Fex J. Structural basis for mechanical transduction in the frog vestibular sensory apparatus: I. The otolithic membrane. Hear Res 45: 179– 190, 1990. doi: 10.1016/S0378-5955(99)00007-6.

29. Cotanche DA. Regeneration of the tectorial membrane in the chick cochlea following severe acoustic trauma. Hear Res 30: 197–206, 1987. doi: 10.1016/0378-5955(87)90136-5.

30. Bielak K, Benkowska-Biernacka D, Ptak M, Stolarski J, Kalka M, Ożyhar A, Dobryszycki P. Otolin-1, an otolith- and otoconia-related protein, controls calcium carbonate bioinspired mineralization. Biochim Biophys Acta - Gen Subj 1867: 1–8, 2023. doi: 10.1016/j.bbagen.2023.130327.

31. Murayama E, Takagi Y, Ohira T, Davis JG, Greene MI, Nagasawa H. Fish otolith contains a unique structural protein, otolin-1. Eur J Biochem 269: 688–696, 2002. doi: 10.1046/j.0014-2956.2001.02701.x.

32. Murayama E, Herbomel P, Kawakami A, Takeda H, Nagasawa H. Otolith matrix proteins OMP-1 and Otolin-1 are necessary for normal otolith growth and their correct anchoring onto the sensory maculae. Mech Dev 122: 791–803, 2005. doi: 10.1016/j.mod.2005.03.002.

33. Beier M, Anken RH, Rahmann H. Calcium-tracers disclose the site of biomineralization in inner ear otoliths of fish. Adv Sp Res 33: 1401–1405, 2004. doi: 10.1016/j.asr.2003.09.044.

34. Ibsch M, Anken R, Beier M, Rahmann H. Endolymphatic calcium supply for fish otolith growth takes place via the proximal portion of the otocyst. Cell Tissue Res 317: 333–336, 2004. doi: 10.1007/s00441-004-0930-6.

35. Ibsch M, Anken RH, Rahmann H. Calcium gradients in the fish inner ear sensory epithelium and otolithic membrane visualized by energy filtering transmission electron microscopy (EFTEM). Adv Sp Res 33: 1395–1400, 2004. doi: 10.1016/j.asr.2003.09.043.

36. Takagi Y. Meshwork arrangement of mitochondria-rich, Na+,K+-ATPase-rich cells in the saccular epithelium of rainbow trout (Oncorhynchus mykiss) inner ear. Anat Rec 248: 483–489, 1997. doi: 10.1002/(SICI)1097-0185(199708)248:4<483::AID-AR1>3.0.CO;2-N.

37. Kwan GT, Smith TR, Tresguerres M. Immunological characterization of two types of ionocytes in the inner ear epithelium of Pacific Chub Mackerel (Scomber japonicus). J Comp Physiol B 190: 419–431, 2020. doi: 10.1007/s00360-020-01276-3.

38. Mayer-Gostan N, Kossmann H, Watrin A, Payan P, Boeuf G. Distribution of ionocytes in the saccular epithelium of the inner ear of two teleosts (Oncorhynchus mykiss and Scophthalmus maximus). Cell Tissue Res 289: 53– 61, 1997. doi: 10.1007/s004410050851.

39. Payan P, Kossmann H, Watrin a, Mayer-Gostan N, Boeuf G. Ionic composition of endolymph in teleosts: origin and importance of endolymph alkalinity. J Exp Biol 200: 1905–1912, 1997.

40. Payan P, Edeyer A, de Pontual H, Borelli G, Boeuf G, Mayer-Gostan N. Chemical composition of saccular endolymph and otolith in fish inner ear: lack of spatial uniformity. Am J Physiol 277: R123–R131, 1999.

41. Tohse H, Murayama E, Ohira T, Takagi Y, Nagasawa H. Localization and diurnal variations of carbonic anhydrase mRNA expression in the inner ear of the rainbow trout Oncorhynchus mykiss. Comp Biochem Physiol B Biochem Mol Biol 145: 257–64, 2006. doi: 10.1016/j.cbpb.2006.06.011.

42. Tohse H, Ando H, Mugiya Y. Biochemical properties and immunohistochemical localization of carbonic anhydrase in the sacculus of the inner ear in the salmon Oncorhynchus masou. Comp Biochem Physiol - A Mol Integr Physiol 137: 87–94, 2004. doi: 10.1016/S1095-6433(03)00272-1.

43. Murayama E, Takagi Y, Nagasawa H. Immunohistochemical localization of two otolith matrix proteins in the otolith and inner ear of the rainbow trout, Oncorhynchus mykiss: Comparative aspects between the adult inner ear and embryonic otocysts. Histochem Cell Biol 121: 155–166, 2004. doi: 10.1007/s00418-003-0605-5.

44. Payan P, Borelli G, Priouzeau F, De Pontual H, Boeuf G, Mayer-Gostan N. Otolith growth in trout Oncorhynchus mykiss: supply of Ca2+ and Sr2+ to the saccular endolymph. J Exp Biol 205: 2687–2695, 2002. doi: Unsp Jeb4023.

45. Grosell M. CO2 and Calcification Processes in Fish. In: Fish Physiology. Academic Press, 2019, p. 133–159.

46. Checkley DM, Dickson AG, Takahashi M, Radich JA, Eisenkolb N, Asch R. Elevated CO2 enhances otolith growth in young fish. Science 324: 1683, 2009. doi: 10.1126/science.1169806.

47. Lombarte A, Lleonart J. Otolith size changes related with body growth, habitat depth and temperature. Environ Biol Fishes 37: 297–306, 1993. doi: 10.1007/BF00004637.

48. Loeppky AR, Belding LD, Quijada-Rodriguez AR, Morgan JD, Pracheil BM, Chakoumakos BC, Anderson WG. Influence of ontogenetic development, temperature, and pCO2 on otolith calcium carbonate polymorph composition in sturgeons. Sci Rep 11: 13878, 2021. doi: 10.1038/s41598-021-93197-6.

49. Mugiya Y, Uchimura T. Otolith resoprtion in duced by anaerobic stress in the goldfish, Carassius auratus. J Fish Biol 35: 813–818, 1989. doi: 10.1111/j.1095-8649.1989.tb03032.x.

50. Meekan MG, Wellington GM, Axe L. El Niño-Southern Oscillation events produce checks in the otoliths of coral reef fishes in the Galápagos Archipelago. Bull Mar Sci 64: 383–390, 1999.

51. Hamilton SL, Kashef NS, Stafford DM, Mattiasen EG, Kapphahn LA, Logan CA, Bjorkstedt EP, Sogard SM. Ocean acidification and hypoxia can have opposite effects on rockfish otolith growth. J Exp Mar Bio Ecol 521, 2019. doi: 10.1016/j.jembe.2019.151245.

52. Andrews AH, Tracey DM, Dunn MR. Lead-radium dating of orange roughy (Hoplostethus atlanticus): Validation of a centenarian life span. Can J Fish Aquat Sci 66: 1130–1140, 2009. doi: 10.1139/F09-059.

53. Beamish RJ, McFarlane GA. A discussion of the importance of aging errors, and an application to walleye pollock: the world’s largest fishery. In: Recent develops in fish otolith research. Columbia: University of South Carolina Press, 1995, p. 545–565.

54. Takagi Y. Otolith formation and endolymph chemistry: A strong correlation between the aragonite saturation state and pH in the endolymph of the trout otolith organ. Mar Ecol Prog Ser 231: 237–245, 2002. doi: 10.3354/meps231237.

55. Takagi Y, Tohse H, Murayama E, Ohira T, Nagasawa H. Diel changes in endolymph aragonite saturation rate and mRNA expression of otolith matrix proteins in the trout otolith organ. Mar Ecol Prog Ser 294: 249–256, 2005. doi: 10.3354/meps294249.

56. Becerra M, Anadon R. Fine structure and development of ionocyte areas in the labyrinth of the trout (Salmo trutta fario). J Anat 183: 463–474, 1993.

57. Lebovitz RM, Takeyasu K, Fambrough DM. Molecular characterization and expression of the (Na+ + K+)-ATPase alpha-subunit in Drosophila melanogaster. EMBO J 8: 193–202, 1989.

58. Lytle C, Xu JC, Biemesderfer D, Forbush B. Distribution and diversity of Na-K-Cl cotransport proteins: a study with monoclonal antibodies. Am J Physiol 269: C1496–C1505, 1995.

59. Abbas L, Whitfield TT. Nkcc1 (Slc12a2) is required for the regulation of endolymph volume in the otic vesicle and swim bladder volume in the zebrafish larva. Development 136: 2837–2848, 2009. doi: 10.1242/dev.034215.

60. Bradford MM. A rapid and sensitive method for the quantitation of microgram quantities of protein utilizing the principle of protein-dye binding. Anal Biochem 72: 248–254, 1976. doi: 10.1016/0003-2697(76)90527-3.

61. Kwan GT, Frable BW, Thompson AR, Tresguerres M. Optimizing immunostaining of archival fish samples to enhance museum collection potential. Acta Histochem 124: 151952, 2022. doi: 10.1016/j.acthis.2022.151952.

62. R Development Core Team. R: A language and environment for statistical computing. R foundation for statistical computing [Online]. 2013. http://www.r-project.org.

63. Köppl C, Wilms V, Russell IJ, Nothwang HG. Evolution of Endolymph Secretion and Endolymphatic Potential Generation in the Vertebrate Inner Ear. Brain Behav Evol 92: 1–31, 2018. doi: 10.1159/000494050.

64. Vavassori S, Mayer A. A new life for an old pump: V-ATPase and neurotransmitter release. J Cell Biol 205: 7–9, 2014. doi: 10.1083/jcb.201403040.

65. Jedrychowska J, Gasanov E V., Korzh V. Kcnb1 plays a role in development of the inner ear. Dev Biol 471: 65–75, 2021. doi: 10.1016/j.ydbio.2020.12.007.

66. Wilms V, Söffgen C, Nothwang HG. Differences in molecular mechanisms of K+ clearance in the auditory sensory epithelium of birds & mammals. J Exp Biol 220: 2701–2705, 2017. doi: 10.1242/jeb.158030.

67. Lim DJ. Ultrastructure of the otolithic membrane and the cupula. A scanning electron microscopic observation. Adv Otorhinolaryngol 19: 35–49, 1973.

68. Tohse H, Saruwatari K, Kogure T, Nagasawa H, Takagi Y. Control of polymorphism and morphology of calcium carbonate crystals by a matrix protein aggregate in fish otoliths. Cryst Growth Des 9: 4897–4901, 2009. doi: 10.1021/cg9006857.

69. Söllner C, Burghammer M, Busch-Nentwich E, Berger J, Schwarz H, Riekel C, Nicolson T. Control of crystal size and lattice formation by starmaker in otolith biomineratization. Science (80-) 302: 282–286, 2003. doi: 10.1126/science.1088443.

70. Kapłon TM, Rymarczyk G, Nocula-Ługowska M, Jakób M, Kochman M, Lisowski M, Szewczuk Z, Ozyhar A. Starmaker exhibits properties of an intrinsically disordered protein. Biomacromolecules 9: 2118–2125, 2008. doi: 10.1021/bm800135m.

71. Weigele J, Franz-odendaal TA. Not All Inner Ears are the Same : Otolith Matrix Proteins in the Inner Ear of Sub-Adult Cichlid Fish, Oreochromis Mossambicus, Reveal Insights Into the Biomineralization Process. Anat Rec 299: 234–245, 2016. doi: 10.1002/ar.23289.

72. Wangemann P, Liu J, Marcus DC. Ion transport mechanisms responsible for K+ secretion and the transepithelial voltage across marginal cells of stria vascularis in vitro. Hear Res 84: 19–29, 1995. doi: 10.1016/0378-5955(95)00009-S.

73. Lang F, Vallon V, Knipper M, Wangemann P. Functional significance of channels and transporters expressed in the inner ear and kidney. Am J Physiol -Cell Physiol 293, 2007. doi: 10.1152/ajpcell.00024.2007.

74. Kimura RS. Distribution, structure, and function of dark cells in the vestibular laby-rinth. Ann Otol Rhinol Laryngol 78: 542–561, 1969. doi: 10.1177/00034894690780031.

75. Ben-Yosef T, Belyantseva IA, Saunders TL, Hughes ED, Kawamoto K, Van Itallie CM, Beyer LA, Halsey K, Gardner DJ, Wilcox ER, Rasmussen J, Anderson JM, Dolan DF, Forge A, Raphael Y, Camper SA, Friedman TB. Claudin 14 knockout mice, a model for autosomal recessive deafness DFNB29, are deaf due to cochlear hair cell degeneration. Hum Mol Genet 12: 2049–2061, 2003. doi: 10.1093/hmg/ddg210.

76. Kitajiri SI, Furuse M, Morita K, Saishin-Kiuchi Y, Kido H, Ito J, Tsukita S. Expression patterns of claudins, tight junction adhesion molecules, in the inner ear. Hear Res 187: 25–34, 2004. doi: 10.1016/S0378-5955(03)00338-1.

77. Roa JN, Munévar CL, Tresguerres M. Feeding induces translocation of vacuolar proton ATPase and pendrin to the membrane of leopard shark (Triakis semifasciata) mitochondrion-rich gill cells. Comp Biochem Physiol -Part A Mol Integr Physiol 174: 29–37, 2014. doi: 10.1016/j.cbpa.2014.04.003.

78. Tresguerres M, Parks SK, Salazar E, Levin LR, Goss GG, Buck J. Bicarbonate-sensing soluble adenylyl cyclase is an essential sensor for acid/base homeostasis. Proc Natl Acad Sci U S A 107: 442–447, 2010. doi: 10.1073/pnas.0911790107.

79. Roa JN, Tresguerres M. Soluble adenylyl cyclase is an acid-base sensor in epithelial base-secreting cells. Am J Physiol - Cell Physiol 311: C340–C349, 2016. doi: 10.1152/ajpcell.00089.2016.

80. Hu MY, Petersen I, Chang WW, Blurton C, Stumpp M. Cellular bicarbonate accumulation and vesicular proton transport promote calcification in the sea urchin larva. Proceedings Biol Sci 287: 20201506, 2020. doi: 10.1098/rspb.2020.1506.

81. Ibsch M, Vöhringer P, Anken RH, Rahmann H. Electronmicroscopic investigations on the role of vesicle-like bodies in inner ear maculae for fish otolith growth. Adv Sp Res 25: 2031–2034, 2000. doi: 10.1016/S0273-1177(99)01011-X.

82. Gauldie RW, Nelson DGA. Aragonite twinning and neuroprotein secretion are the cause of daily growth rings in fish otoliths. Comp Biochem Physiol -- Part A Physiol 90: 501–509, 1988. doi: 10.1016/0300-9629(88)90227-7.

83. Kwan GT. Characterization of Marine Teleost Ionocytes in the Gill, Skin, and Inner Ear Epithelia and their Implications for Ocean Acidification. University of California, San Diego: 2020.

84. Mass T, Giuffre AJ, Sun C, Stifler CA, Frazier MJ, Neder M, Tamura N, Stan C V., Marcus MA, Gilbert PUPA. Amorphous calcium carbonate particles form coral skeletons. Proc Natl Acad Sci 114: E7670–E7678, 2017. doi: 10.1073/pnas.1707890114.

85. Barron ME, Thies AB, Espinoza JA, Barott KL, Hamdoun A, Tresguerres M. A vesicular Na+/Ca2+ exchanger in coral calcifying cells. PLoS One 13: e0205367, 2018. doi: 10.1371/journal.pone.0205367.

86. Mummadisetti MP, Drake JL, Falkowski PG. The spatial network of skeletal proteins in a stony coral. J R Soc Interface 18: 20200859, 2021. doi: 10.1098/rsif.2020.0859.

87. Nakaya K, Harbidge DG, Wangemann P, Schultz BD, Green ED, Wall SM, Marcus DC. Lack of pendrin HCO-3 transport elevates vestibular endolymphatic [Ca2+] by inhibition of acid-sensitive TRPV5 and TRPV6 channels. Am J Physiol - Ren Physiol 292: 1314–1321, 2007. doi: 10.1152/ajprenal.00432.2006.

88. Wangemann P, Nakaya K, Wu T, Maganti RJ, Itza EM, Sanneman JD, Harbidge DG, Billings S, Marcus DC. Loss of cochlear HCO3-secretion causes deafness via endolymphatic acidification and inhibition of Ca2+ reabsorption in a Pendred syndrome mouse model. Am J Physiol - Ren Physiol 292: 1345–1353, 2007. doi: 10.1152/ajprenal.00487.2006.

89. Takagi Y. Ultrastructural immunolocalization of the otolith water-soluble-matrix in the inner ear of rainbow trout just-hatched fry. Fish Sci 66: 71–77, 2000. doi: 10.1046/j.1444-2906.2000.00010.x.

90. Mitrovic D, Dymowska A, Nilsson GE, Perry SF. Physiological consequences of gill remodeling in goldfish (Carassius auratus) during exposure to long-term hypoxia. AJP Regul Integr Comp Physiol 297: R224–R234, 2009. doi: 10.1152/ajpregu.00189.2009.

91. Phuong LM, Huong DTT, Nyengaard JR, Bayley M. Gill remodelling and growth rate of striped catfish Pangasianodon hypophthalmus under impacts of hypoxia and temperature. Comp Biochem Physiol Part A Mol Integr Physiol 203: 288–296, 2017. doi: 10.1016/j.cbpa.2016.10.006.

92. Matey V, Richards JG, Wang Y, Wood CM, Rogers J, Davies R, Murray BW, Chen X-Q, Du J, Brauner CJ. The effect of hypoxia on gill morphology and ionoregulatory status in the Lake Qinghai scaleless carp, Gymnocypris przewalskii. J Exp Biol 211: 1063–1074, 2008. doi: 10.1242/jeb.010181.

93. Navarro MO, Kwan GT, Batalov O, Choi CY, Pierce NT, Levin LA. Development of Embryonic Market Squid, Doryteuthis opalescens, under Chronic Exposure to Low Environmental pH and [O2]. PLoS One 11: e0167461, 2016. doi: 10.1371/journal.pone.0167461.

94. Passerotti MS, Jones CM, Swanson CE, Quattro JM. Fourier-transform near infrared spectroscopy (FT-NIRS) rapidly and non-destructively predicts daily age and growth in otoliths of juvenile red snapper Lutjanus campechanus (Poey, 1860). Fish Res 223: 105439, 2020. doi: 10.1016/j.fishres.2019.105439.

95. Helser TE, Benson I, Erickson J, Healy J, Kastelle C, Short JA. A transformative approach to ageing fish otoliths using fourier transform near infrared spectroscopy: A case study of eastern bering sea walleye pollock (gadus chalcogrammus). Can J Fish Aquat Sci 76: 780–789, 2019. doi: 10.1139/cjfas-2018-0112.

96. Passerotti MS, Helser TE, Benson IM, Barnett BK, Ballenger JC, Bubley WJ, Reichert MJM, Quattro JM. Age estimation of red snapper (Lutjanus campechanus) using FT-NIR spectroscopy: feasibility of application to production ageing for management..

97. Healy J, Helser TE, Benson IM, Tornabene L. Aging Pacific cod (Gadus macrocephalus) from otoliths using Fourier-transformed near-infrared spectroscopy . Ecosphere 12, 2021. doi: 10.1002/ecs2.3697.

